# Targeting monocytic Occludin impairs monocyte transmigration and HIV neuroinvasion

**DOI:** 10.1101/2023.09.11.557242

**Authors:** Diana Brychka, Nilda Vanesa Ayala-Nunez, Yonis Bare, Amandine Dupas, Emma Partiot, Vincent Mittelheisser, Vincent Lucansky, Jacky G. Goetz, Nael Osmani, Raphael Gaudin

## Abstract

Transmigration of circulating monocytes from the bloodstream toward the central nervous system (CNS) represents a hallmark of neuroinflammation and plays an important role during viral encephalitis and HIV-associated neurocognitive disorders (HAND). The molecular mechanisms involved in monocyte transmigration through endothelia has been extensively studied, but how monocytes locally unzip tight junction-associated proteins (TJAPs) of the endothelium composing the neurovascular unit (NVU) to reach the CNS remains poorly understood. Here, we show that human circulating monocytes express the TJAP Occludin (OCLN) to promote transmigration through cerebral microvessel endothelial cells. Silencing monocytic OCLN (mOCLN) impairs monocyte transmigration, while mOCLN overexpression increases transmigration. Using high-resolution live cell imaging, we observed that mOCLN clusters at the monocyte-endothelium interface during the transmigration process, forming a transient ring of mOCLN at the site of diapedesis. Furthermore, we designed OCLN-derived peptides targeting its extracellular loop (EL) 1 or 2 to prevent potential trans-homotypic interactions of mOCLN with endothelial OCLN. We found that transmigration of human monocytes was significantly inhibited upon treatment with the EL2 peptide *in vitro* and in zebrafish embryos, while preserving vascular integrity. Monocyte transmigration toward the brain is an important process for HIV neuroinvasion and here, we showed that the treatment of transmigrating monocytes with the EL2 peptide prevents the dissemination of HIV to cerebral organoids. In conclusion, our study identifies an important role for monocytic OCLN during transmigration and provides a proof-of-concept for the development of mitigation strategies to prevent HIV neuroinvasion.

## Introduction

The neurovascular unit (NVU), also referred to as the blood-brain barrier (BBB), is a complex structure, which integrity is critical to protect the central nervous system (CNS). Passage through the NVU is a tightly regulated process, and monocyte infiltration represents a hallmark of CNS inflammation. Indeed, monocytes are the first immune cells recruited to the CNS and a major contributor of neurological disorders, particularly important in the context of viral encephalitis (1, 2). The transmigration of immune cells from the bloodstream toward the CNS is mediated by their adherence to the endothelial cell wall of the NVU, followed by squeezing of the immune cell in between endothelial cells, and release in the CNS. The ability of monocytes to transmigrate through the NVU is a pioneering event that can tilt the fragile balance between a healthy or a disordered brain. Understanding the molecular mechanisms involved in monocyte transmigration to the brain is thus highly needed to envision novel strategies to modulate neuroinvasion of immune cells and inhibit neurotropic infections.

Monocyte transmigration is a multistep process requiring a number of essential molecular partners at the cell surface as well as dramatic cytoskeleton and membrane rearrangements to cross an endothelia (3). The NVU exhibits an additional level of complexity, as it possesses tight junctions (TJs), which make the NVU the most impermeant endothelia of the human body. However, the mechanisms used by monocytes to cross endothelial TJs during transmigration, while avoiding NVU disruption, remains incompletely understood. Endothelial cells composing the NVU are formed of specific TJ-associated proteins (TJAPs), including Claudine-5 (CLDN5) and Occludin (OCLN). These proteins have been designated as potential targets for the prevention of viral neuroinvasion (4). Hence, a better understanding of the molecular mechanism involved in monocyte transmigration is important to develop new antiviral strategies that would aim at preventing neuroinvasion.

The CNS is considered a major HIV reservoir, which infection causes HIV-associated neurocognitive disorders (HAND) (5). It was previously proposed that the human immunodeficiency virus (HIV) is taking advantage of blood-circulating monocytes, enhancing transmigration to cross the NVU using a “Trojan horse” strategy (6, 7). Infected monocyte-derived macrophages have been detected in the brain of patients despite antiretroviral therapy (ART) (8), but how and when they have been infected remains mostly obscure. Monocytes are able to capture and store infectious viral particles without being infected themselves (9), a “hiding” strategy that could play a major role during HIV dissemination to the CNS. Although infrequent, several reports from different groups also indicated that circulating monocytes contain integrated HIV-1 DNA, even in individuals under ART (10–13), highlighting the importance of monocytes as a potential source of infected-macrophages in the CNS. Therefore, a better understanding of the molecular mechanisms leading to monocyte transmigration is crucial to envision novel strategies to prevent HIV neuroinvasion.

Several TJAPs have been proposed to play a role during leukocyte transmigration (3, 14) and here, we reinvestigated the function of the most expressed monocytic TJAPs (mTJAPs) during this process. First, we performed a mini-screen in THP-1 monocytic cells using siRNA targeting the most expressed mTJAP RNAs. OCLN silencing in THP-1 cells significantly decreases the ability of monocytes to transmigrate through the human cerebral microvascular endothelial D3 cell line (hCMEC/D3), while mOCLN overexpression in human primary monocytes, isolated from healthy blood donors, results in increased transmigration. Furthermore, we find that monocytic OCLN (mOCLN) accumulates transiently at the interface between monocytes and endothelial cells. To prevent interaction between mOCLN and endothelial OCLN, we designed short peptides copying regions of the extracellular loop 1 or 2 of OCLN (EL1 and EL2 respectively). We show that the EL2 peptide significantly prevents monocyte transmigration *in vitro* and in zebrafish, while preserving endothelial impermeability. Finally, we reveal that treatment with the OCLN-derived peptides prevents monocyte infiltration and HIV replication in cortical organoids. In conclusion, our work highlights the importance of mOCLN during monocyte transmigration and proposes a strategy based on the use of the OCLN-derived EL2 peptide to control the access of monocytes to the CNS, offering innovative mitigation strategies in the context of HIV brain reservoir establishment, although also opening other applications.

## Results

### Monocytic Occludin promotes monocyte transmigration through BBB-like endothelia

To evaluate whether tight-junction associated proteins (TJAPs) are expressed by monocytes, we first interrogated RNAseq databases (15), seeking for TJAP-coding mRNAs in human primary monocytes and the monocytic cell line THP-1 (**Figure 1A**). We found that a subset of TJAP-coding mRNAs was expressed by these cells. To test if the monocytic expression of TJAPs could participate in monocyte transmigration, we knocked-down the 21 most expressed mTJAP-coding RNAs in THP-1 cells using siRNA and performed a transmigration assay through cerebral microvascular endothelial cells clone D3 (hCMEC/D3) (16) (**Figure 1B**). Compared to control, silencing of OCLN resulted in a two-fold decrease of the number of transmigrated monocytes (**Figure 1C**), suggesting that monocytic OCLN (mOCLN) may participate in monocyte transmigration through BBB-like endothelia. To further confirm this observation, we built a lentivector expressing 2 guide RNAs (gRNAs) targeting OCLN, a Cas9-specific tracrRNA and the Cas9 enzyme (**Figure 1D**, and Addgene #208398). This construct also codes for puromycin resistance and the E2-Crimson far-red fluorescent protein to help for the selection of transduced cells. This construct was used to produce lentivectors and transduce THP-1 cells, allowing to obtain a CRISPR edited THP-1 OCLN knock out (KO) cell line. Western blot analysis of OCLN protein expression in these cells showed no band at expected size (**Suppl Figure S1A**). Of note, a band of very dim intensity and at a lower size was observed, which nature is unknown, but whether this represents some remaining OCLN isoforms or non-specific staining, its intensity remains at marginal levels compared to control cells. This cell line was the only clone of THP-1 that showed clear OCLN disappearance. In order to thereafter compare transmigration from the same clonal cells, THP-1 OCLN KO cells were rescue using an EGFP-OCLN coding vector or an EGFP-OCLN with deletion of the C-terminal tail (EGFP-OCLN-ΔC). As a control, THP-1 OCLN KO cells were transduced with an irrelevant EGFP-CAAX coding vector, CAAX being the farnelysation motif of the HRas protein known to address it to the plasma membrane (PM; (17)). Transmigration of THP-1 EGFP-OCLN showed significantly more transmigration potency than their EGFP-CAAX and EGFP-OCLN-ΔC counterparts (**Figure 1E**), further confirming the importance of OCLN during monocyte transmigration.

**Figure 1.**
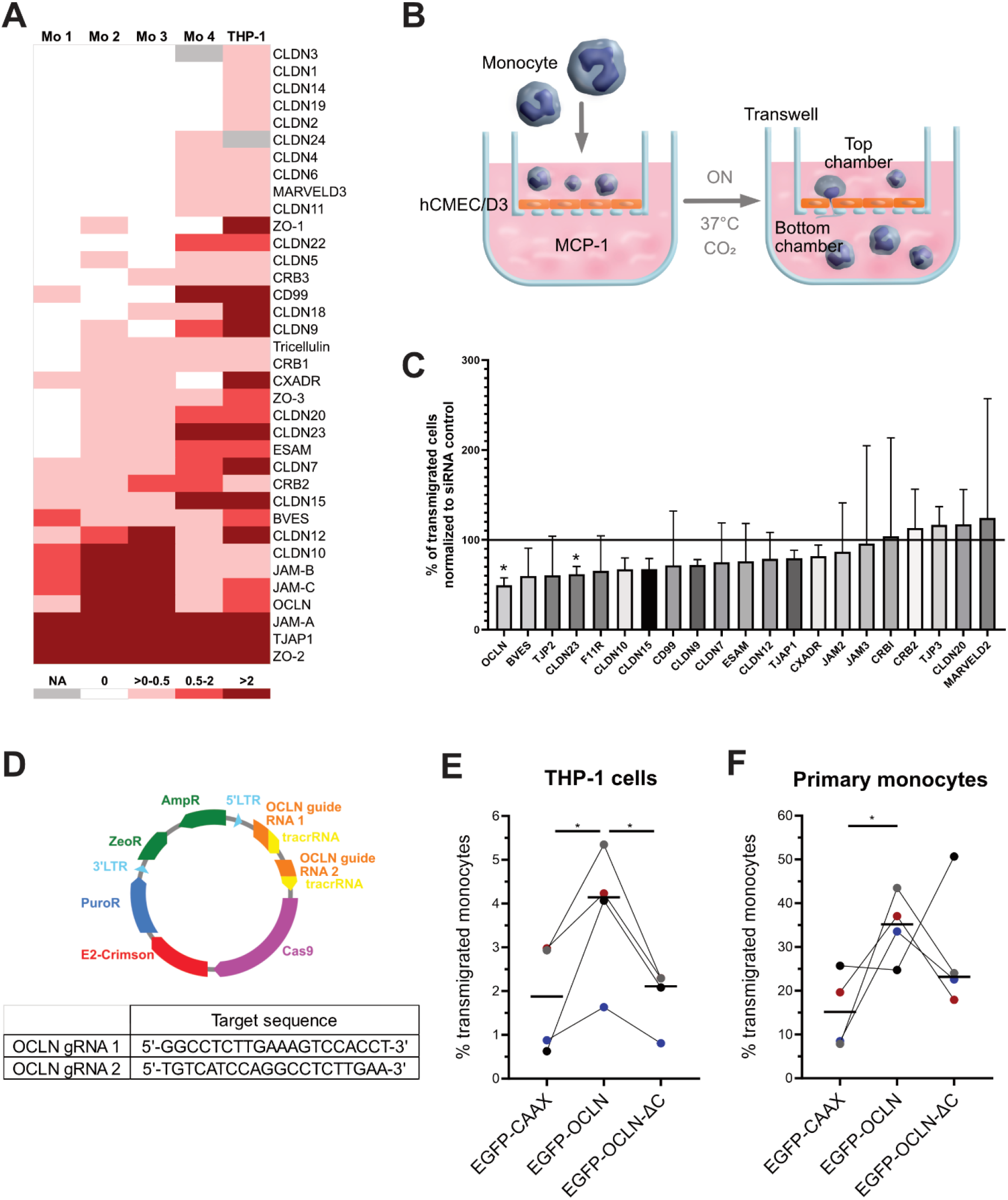
Monocytic OCLN favors monocyte transmigration through BBB-like endothelial cells. (**A**) Heatmap of TJAP-coding mRNAs in human primary monocytes and the THP-1 cell line based on existing RNAseq databases (see Material & Methods). (**B**) Schematic of the experimental setup of a transmigration assay with monocytes crossing a confluent monolayer of brain endothelial cells (hCMEC/D3). Either primary human monocytes or THP-1 cells are added to the top chamber and transmigration is allowed to happen 17 h (overnight) at 37°C, 5% CO_2_. (**C**) An siRNA screen of TJAP proteins was performed to define the role of these proteins on monocyte transmigration. The TJAPs were selected based on the heatmap shown in A. THP-1 cells were electroporated with the indicated siRNAs and a transmigration assay was done. The data is presented as a percentage of transmigrated THP-1 cells normalized to the siRNA control (black line). The graph shows the mean +/-SD of four independent experiments. Two-tailed Student T test shows p value < 0.05 (*). (**D**) Schematic representation of a CRISPR/Cas9 lentiviral construct used for the generation of OCLN KO THP-1 cells with indicated target sequences. (**E**) Percentage of transmigrated OCLN KO THP-1 cells rescued either with EGFP-CAAX, EGFP-OCLN, or EGFP-ΔC-OCLN across hCMEC/D3 monolayer. The data represents the mean from four individual experiments. Two-tailed Student T test shows p value < 0.05 (*). (**F**) Percentage of transmigrated primary GFP+ monocytes transduced either with EGFP-CAAX, EGFP-OCLN, or EGFP-ΔC-OCLN across hCMEC/D3 monolayer. The data represents the mean values obtained from two donors in duplicates. Two-tailed Student T test shows p value < 0.05 (*).

In primary human monocytes, endogenous mOCLN was readily observed by western blot and upon OCLN overexpression (**Suppl. Figure S1B-C**). Despite many attempts however, we could not detect the endogenous OCLN expressed by monocytes by immunofluorescence (IF), likely because it is weakly expressed, exhibiting very low signal-to-noise ratio. To further evaluate the role of OCLN in primary cells during transmigration, we optimized a protocol for efficient human monocyte transduction using a Vpx-based strategy (18). We overexpressed EGFP, EGFP-OCLN or EGFP-OCLN-ΔC in primary monocytes, which showed variable efficiency depending on the construct (**Suppl. Figure S1D**). Transduction was > 10% efficiency in monocytes, which is acceptable for primary cell transduction and would not majorly influence the following experiments as we selected only the fluorescent (transduced) cells. Analysis of the percentage of transduced primary monocytes that transmigrated over the ones that did not transmigrate revealed that overexpression of EGFP-OCLN significantly increased primary monocyte transmigration through hCMEC/D3, while the OCLN-ΔC mutant did not (**Figure 1F**). Of note, our attempts to knock-down/out OCLN in primary monocytes were unsuccessful, which might be due to the relatively stable nature of OCLN, while primary monocytes are kept in culture for only 3 days maximum before they start to significantly differentiate or die.

Together, our data indicates that the expression of OCLN by monocytes is favoring transmigration of monocytes through BBB-like endothelia.

### Distribution of monocytic OCLN during transmigration

To better understand the dynamics of mOCLN distribution during monocyte transmigration through hCMEC/D3 cells, we optimized imaging conditions to support 5D live cell microscopy (X, Y, Z, Time, Channels). Catching transmigration events being rare and the imaging being particularly challenging, we were unable to quantify 5D movies, but could reproducibly see the following events: upon early steps of monocyte attachment to the endothelial monolayer, mOCLN was mostly found at the PM of monocytes, with some OCLN polarizating toward the endothelium side (**Figure 2A-B**, **Movie S1** and **Movie S2**). During the transmigration process, OCLN clusters at the monocyte-endothelium interface, while the monocytes started to protrude across. At this stage, we observed the formation of OCLN-containing internal compartments in the monocytes. The OCLN-containing compartments forming during transmigration were not classical recycling endosomes (Rab11- and Rab13-negative; **Suppl. Figure S2A-B**) nor degradative compartments (Rab7-negative; **Suppl. Figure S2C**). They were truly internal (as opposed to deep PM invagination) as fluorescent Dextran, a fluid phase marker, could not access it (**Suppl. Figure S2D**). Toward the end of the transmigration process, the monocyte’s whole cell body squeezed in between endothelial cells and flattened beneath the endothelium, in close contact with the coverslip. At these later stages, OCLN was distributed at the PM and in clusters that may be internal. One of the most striking observation of the transmigration process is that OCLN transiently relocalized to inter-endothelial junctions (**Figure 2A-B**). In particular, we caught a very transient phenomenon, where an OCLN ring formed at the monocyte-endothelia interface (**Figure 2B** right panels, and **Movie S2**), with has faster kinetics than our imaging time interval (10 min). Such structure was never observed in EGFP-CAAX-expressing monocytes (**Figure 2C** and **Movie S3**). Finally, we observed that monocytes transduced with EGFP-OCLN-ΔC were not protruding through endothelia, although the protein also polarized toward the endothelial monolayer (**Figure 2D** and **Movie S4**), suggesting that the C-terminal domain of mOCLN is required to initiate the formation of protrusions and/or the opening of endothelial cell junctions.

**Figure 2.**
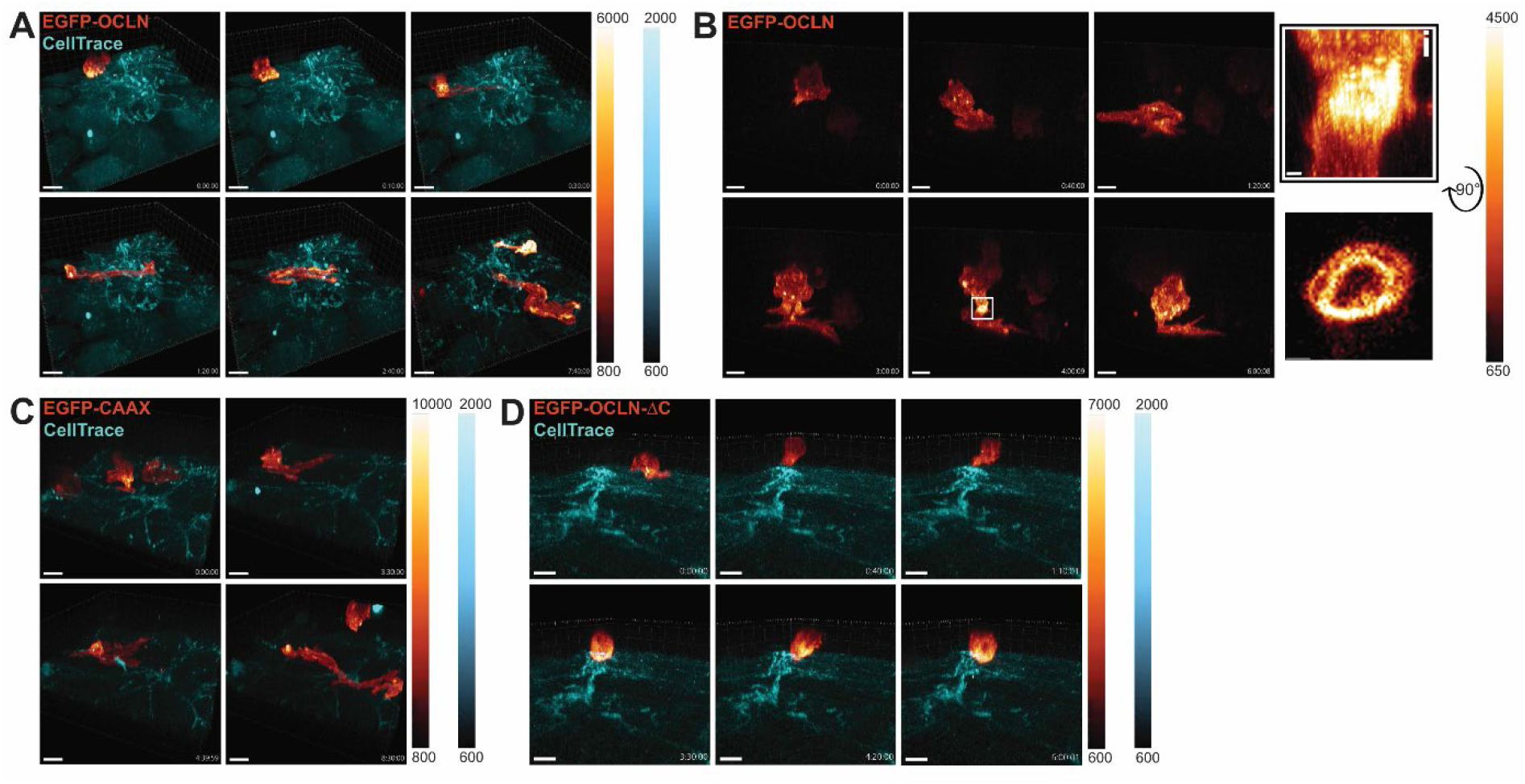
Monocytic OCLN transiently polarizes at monocyte-endothelial interaction sites during transmigration. (**A-D**) 3D time-lapse spinning disk confocal microscopy of transmigrating monocytes. Fluorescence intensity scale is shown on the side of each panel. EGFP is represented using the “red Fire” color-coding, and endothelial cells are stained with a CellTrace marker (cyan) added the addition of the monocytes prior. Timescales are shown is the bottom right corner and corresponds to the time after which imaging acquisition started. (**A**) Imaging of a primary monocyte transduced with EGFP-OCLN on hCMEC/D3 monolayer. Images were taken every 10 min. Scale bar: 10 μm. Full video can be found in **Movie S1**. (**B**) Imaging of a THP-1 cell transduced with EGFP-OCLN on hCMEC/D3 monolayer (not stained). Images were taken every 10 min. Scale bar: 10 μm. Full video can be found in **Movie S2**. The white square highlights OCLN accumulation at the potential interface between hCMEC/D3 cells and the THP-1 cell. (i) Enlarged image from the white square and a 90° flip showing the accumulation of OCLN. Scale bar: 1 µm. (**C**) Imaging of a primary monocyte transduced with EGFP-CAAX on hCMEC/D3 monolayer. Images were taken every 10 min. Scale bar: 10 μm. Full video can be found in **Movie S3**. (**D**) Imaging of a primary monocyte transduced with EGFP-OCLN-ΔC on hCMEC/D3 monolayer. Images were taken every 10 min. Scale bar: 10 μm. Full video can be found in **Movie S4**.

### An OCLN-derived peptide inhibits monocyte transmigration *in vitro*

Peptides derived from the extracellular loops (EL) of OCLN were shown to bind to OCLN, indicating that transcellular homotypic OCLN interactions between cells occur (19–24). In order to prevent the potential interactions of mOCLN with endothelial OCLN, we designed peptides mimicking the extracellular loop 1 (EL1) or 2 (EL2) of the human OCLN and associated scramble controls (scrEL1 and scrEL2 respectively; **Figure 3A**) based on previous literature ((19–24); see Material & Methods for details). None of our peptides significantly changed hCMEC/D3 permeability *in vitro* (**Figure 3B**).

**Figure 3.**
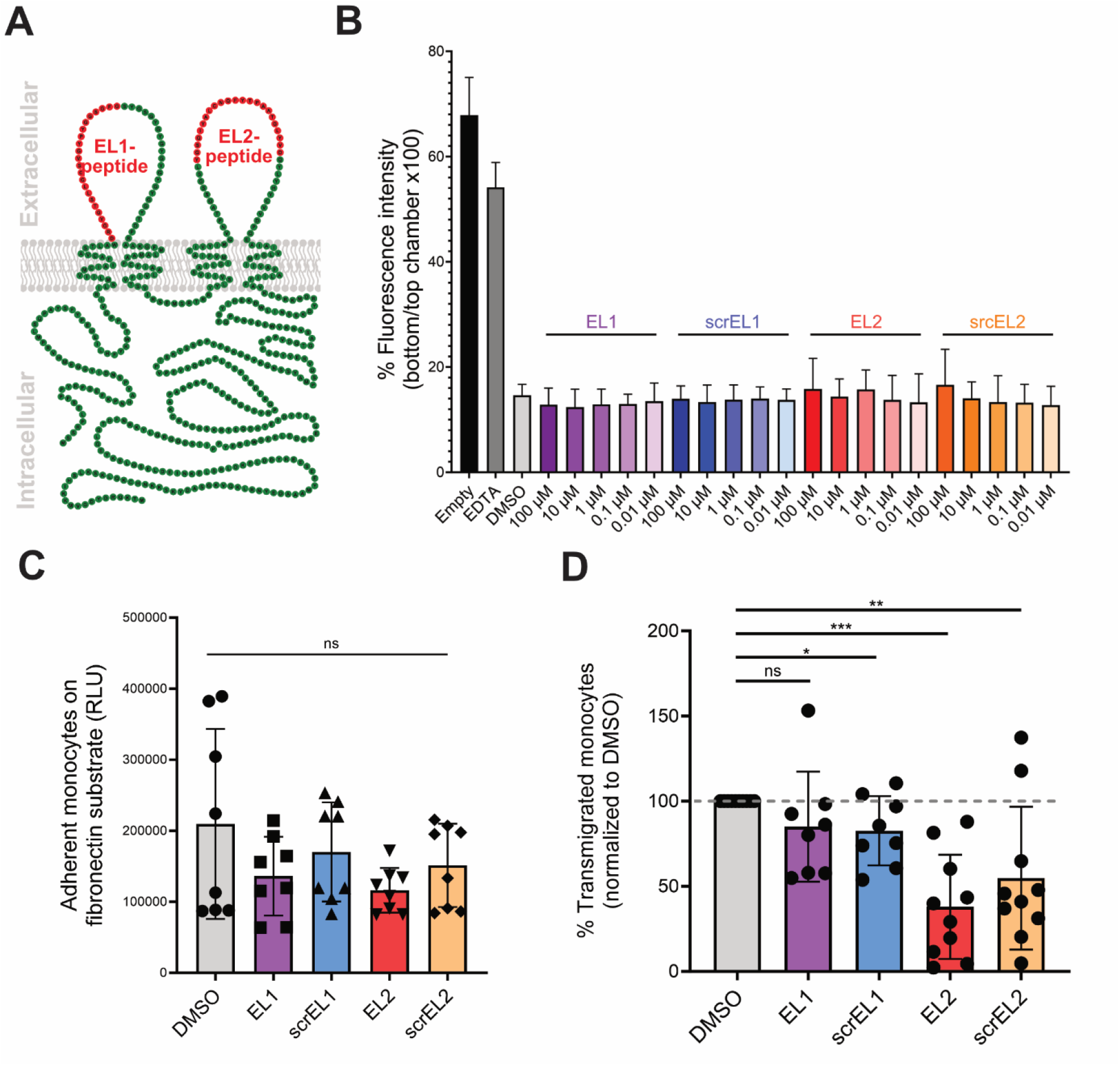
OCLN-derived EL2 peptide inhibits monocyte transmigration through BBB-like endothelial cells. (**A**) Schematic representation of OCLN (green) at the PM with OCLN-derived peptides highlighted in red (see sequence in Material & Methods). (**B**) Permeability of hCMEC/D3 monolayer after overnight exposure of the monolayer to OCLN-derived peptides or their scramble controls. (**C**) Adhesion of primary monocytes on fibronectin after pre-incubation with OCLN-derived peptides for 1 h measured with CellTiter Glo. Adhesion time is 30 min. Each point represents a measurement. n=4 donors. Error bars are SD. There was no statistical difference between all the conditions (ns = non-significant). (**D**) Percentage of primary monocytes after 1 h exposure to OCLN-derived peptides or their scramble controls that transmigrated across the hCMEC/D3 monolayer overnight. Each point represents one donor (average of a duplicate). Error bars are SD. Two-tailed unpaired Student t test shows p value < 0.05 (*), < 0.01 (**), or < 0.001 (***).

We next evaluated the impact of the peptides on the various stages of monocyte transmigration. First, we found that none of the peptides significantly perturbed monocyte adhesion to fibronectin, the main extracellular matrix at the apical surface of hCMED/D3 cells (**Figure 3C**). Strikingly, the treatment of human primary monocytes with the EL2 peptide significantly decreased their transmigration through hCMEC/D3 cells, while EL1 peptide did not (**Figure 3D**). Of note, the scrEL2 peptide also exhibited some inhibitory activity, potentially because negatively charged amino acids of the EL2 sequence were partially conserved.

Together, this data suggests that EL2 peptide could represent an interesting strategy to target OCLN-mediated monocyte transmigration.

### The OCLN-derived EL2 peptide is not toxic while inhibiting monocyte transmigration *in vivo*

We next aimed to evaluate the potency of the OCLN-derived peptides to inhibit human monocyte transmigration. To this end, we took advantage of a method that we previously established (25), consisting of injecting pre-labeled human monocytes in zebrafish embryos expressing a fluorescent endothelium (*Tg(fli1a:egfp)*; (**Figure 4A-B**). This model allows efficient tracking of human monocytes, including virus-infected ones (25), transmigrating across endothelial cells of the vascular caudal plexus. Human OCLN has a 95% consensus sequence with zebrafish OCLN (zOCLN) as calculated upon TCoffee alignment ((26); **Suppl. Figure S3A**). The EL1 and EL2 peptide derived from human OCLN had partial, but significant similarities (**Suppl. Figure S3B**). First, we tested for toxicity and/or vascular permeability in zebrafish embryos by injecting solely the EL1 or EL2 peptides. We did not observe developmental defects as estimated by the morphology and heartbeat of the embryos, and found no significant leakage of the vascular endothelium between the conditions (**Suppl. Figure S3C-F**). As a control, LPS injection induced vascular leakage, as previously described (27). Next, human primary monocytes from four individual blood donors with DMSO, EL1, or EL2 peptides were intravenously (i.v) injected in 48 hpf zebrafish embryos and confocal imaging of the vascular caudal plexus was performed at 6-8 h post-injection (**Figure 4C**). Interestingly, the human OCLN-derived EL2 peptide significantly decreased monocyte transmigration, despite relatively high interdonor-variability (**Figure 4D**), which mirror the data obtained *in vitro*.

**Figure 4.**
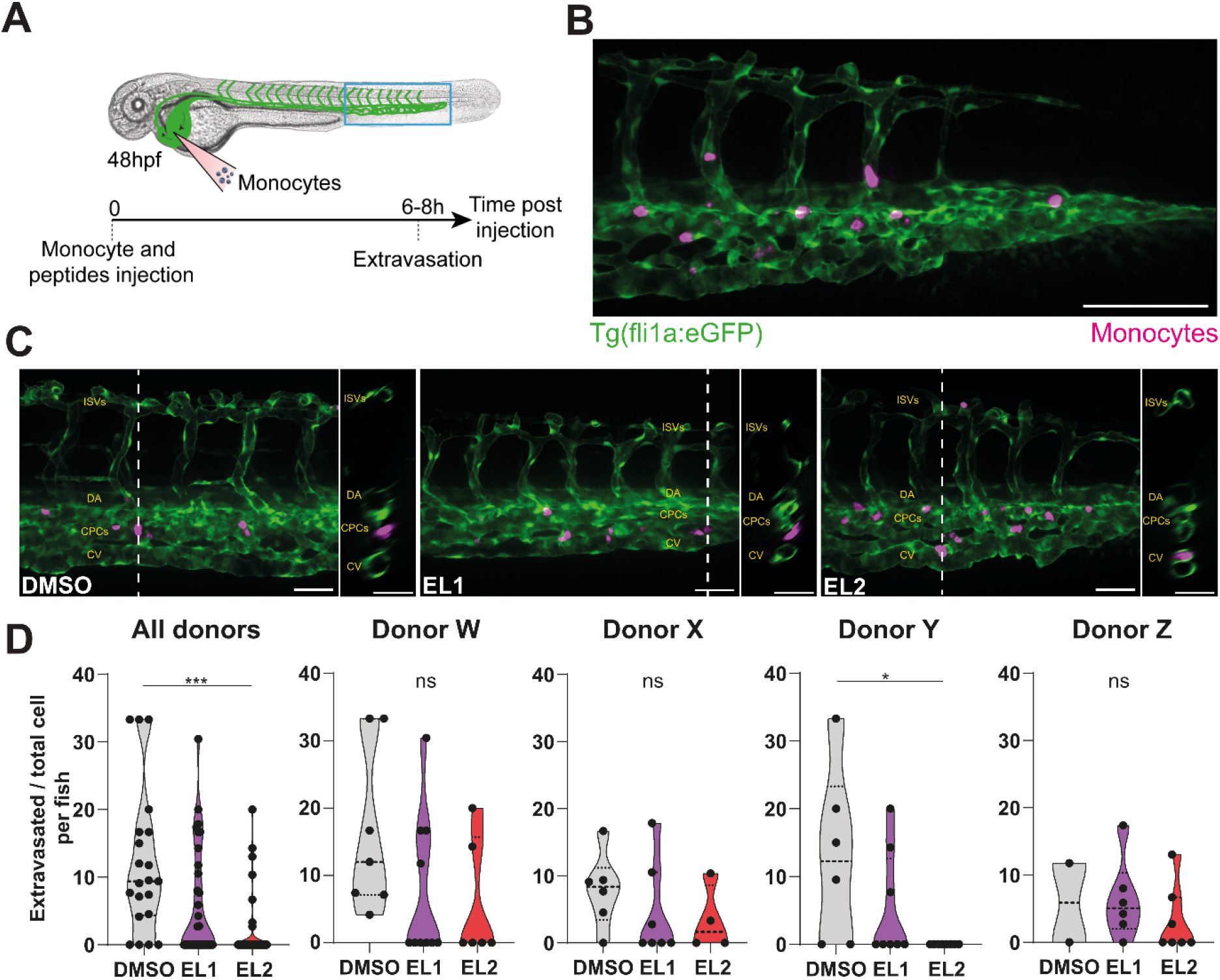
OCLN-derived EL2 peptide inhibits human monocyte transmigration *in vivo*. (A) Representative scheme of the experimental design associated with the zebrafish model. Human primary monocytes were with CellTrace, and injected into the duct of Cuvier of Tg(fli1a:eGFP-CAAX) zebrafish embryos (GFP-labeled endothelial cells). (B) Zebrafish imaging was done at 6–8 h post injection by spinning-disk confocal microscopy. Representative z-projection of confocal images of monocytes from patient X arrested in the vasculature at the tail of a zebrafish embryo at 6 hours post injection Scale bar: 100 µm. (C) Representative images and orthoslice of monocytes at 6 hpi. Scale bar: 50 µm. (D) Violin plots of the ratio of extravasated monocytes from four donors (merged and seperated) at 6–8 h post injection. Each dot represents the ratio calculated from all the monocytes tracked within a fish. The experiment was carried out four independent times by using four different monocyte donors with nDMSO=21, nEL1=31, nEL2=25 in total (nDMSO=7, nEL1=10, nEL2=6 for donor W; nDMSO=6, nEL1=7, nEL2=4 for donor X; nDMSO=6, nEL1=8, nEL2=8 for donor Y; nDMSO=2, nEL1=6, nEL2=7 for donor Z). P value < 0.05 (*), <0.005 (**), and <0.0001 (***). The statistical difference of non-Gaussian datasets was analyzed using the Kruskal-Wallis test with Dunn’s post hoc test.

Together, this data highlights that the EL2 peptide is a potent inhibitor of monocyte transmigration that retain activity *in vivo*, while not exhibiting noticeable toxicity.

### Monocytes carrying HIV-1 particles exhibit increased transmigration

Finally, we wanted to test whether the EL2 peptide could be used in a human pathological context, using HIV-1 as a model system. Indeed, HIV-1 has been shown to associate with monocytes and invades the brain using the “Trojan horse” strategy, consisting of being carried through the BBB by hiding in transmigrating circulating cells (7). Here, we showed in our model that the exposure of primary monocytes to HIV-1 promotes monocyte transmigration (**Suppl. Figure S4A**), as previously reported (28). While monocyte-derived macrophages and microglia are permissive targets allowing HIV-1 productive infection, circulating monocytes are poorly permissive to HIV-1 infection (13). Nevertheless, monocytes can capture and store infectious viral particles without being infected themselves (9). Here, we show that primary monocytes exposed to fluorescent HIV-1 Gag-iGFP R5 particles are highly positive for EGFP, regardless of antiviral AZT treatment (**Suppl. Figure S4B**), indicating that the virus particles are carried by monocytes but do not (or very poorly) replicate in them. At the subcellular level, monocytes exposed to Gag-iGFP fluorescent particles exhibit EGFP puncta positive for anti-Gag p17antibody that remain in AZT-treated monocytes (**Suppl. Figure S4C** middle panels). Using a VSV-G pseudotyped virus to force productive infection, we also observed the dotted structures, but also a diffuse staining, suggestive of neosynthesis of Gag in the cytosol, which was not observed in AZT-treated cells (**Suppl. Figure S4C** right panels).

Together, this data highlights that HIV is carried by circulating monocytes into intracellular structures, while productive infection can be observed by the appearance of a diffuse cytosolic Gag staining.

### The OCLN-derived EL2 peptide prevents monocyte-mediated HIV neuroinvasion

To mimic monocyte-mediated HIV neuroinvasion, we took advantage of the monocyte-infiltrating cortical organoid assay we previously developed in the context of Zika virus neuroinfection (25). Briefly, we differentiated cortical organoids from human embryonic stem (hES) cells and incubated it with human primary monocytes previously exposed to fluorescent HIV-1 (**Figure 5A**). Using long-term imaging, we were able to monitor in live the infiltration of primary monocytes carrying HIV-1 Gag-imCherry particles (29) for 8 days (**Figure 5B** and **Movie S5**). Here, we observed for the first time that monocytes carrying HIV-1 particles can infiltrate the organoid and started to express increased diffuse Gag-imCherry fluorescence (**Figure 5B**, yellow arrows), suggesting that active viral replication was initiated (as in **Suppl. Figure S4C**). At high resolution in fixed samples, we indeed detected infiltrated monocyte-derived cells productively infected by HIV-1 (puncta and cytosolic Gag-imCherry fluorescence) deep into the organoid (**Figure 5C-E**). This data suggests that infiltration/differentiation of monocytes in the organoids is associated with the induction of productive HIV-1 replication. Finally, we placed cortical organoids in the bottom chamber of transwells and added HIV-exposed primary monocytes treated with OCLN-derived peptides in the top chamber to allow transmigration to occur (**Figure 5E**). In this setup, we showed that HIV-infected monocytes allowed the passage of HIV through the BBB regardless of ART treatment (**Figure 5F**, DMSO and DMSO AZT treatment). In contrast, treatment with EL1 and EL2 peptides significantly decreased the amount of HIV-1 RNA that crossed the endothelium (**Figure 5F**).

**Figure 5.**
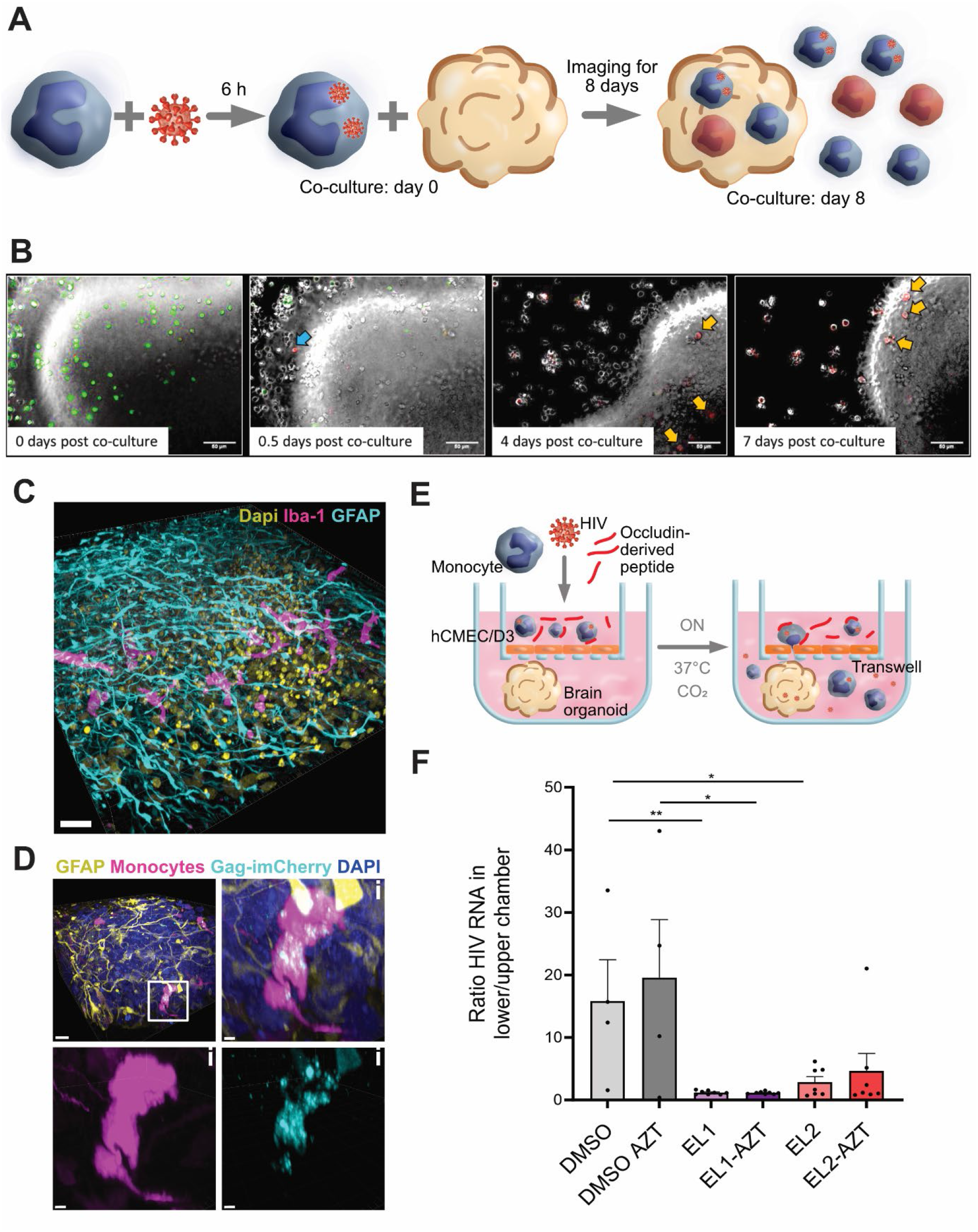
OCLN-derived EL2 peptide inhibits HIV-1 neuroinvasion. (**A**) Schematic of the experimental setup. (**B**) Time-lapse imaging as shown in A of primary human monocytes pre-stained with CellTrace (green) and exposed to HIV-1 Gag-imCherry (red) for 6 h prior to the beginning of the co-incubation of monocytes with a hESC-derived cortical organoid. Images were taken every hour for 8 days. At day 0 of co-incubation, monocytes carrying HIV-1 particles show red dots (not visible at this resolution). At 18 hpi (corresponding to 0.5 days post-co-incubation, upper right panel), very few cells were productively infected. Starting from day 4 up until day 8, multiple cells were observed to be productively infected and penetrating inside the cerebral organoid (highlighted with yellow arrows). Scale bar: 5 μm. Full video can be found in **Movie S5**. (**C**) 3D reconstruction of an immunofluorescent image of a brain organoid with non-infected primary monocytes. Monocyte-derived cells are stained with Iba1 (magenta), astrocytes with GFAP (cyan) and cell nuclei with Dapi (yellow). Scale bar: 40 μm. (**D**) Three-dimensional reconstruction of an immunofluorescence image of a hESC-derived cortical organoid infiltrated with primary monocytes (CellTrace, magenta), infected with HIV-1 Gag-imCherry particles (cyan) for 7 days after the beginning of co-incubation (as in A). Astrocytes are stained with GFAP (yellow) and cell nuclei with Dapi (blue). Scale bar: 15 μm. The infected monocyte highlighted in the white square is magnified in (i). Scale bar: 5 μm. (**E-F**) Transmigration of primary monocytes exposed to HIV-1 R5 Gag-imCherry for 24h, in the presence or absence of 1 µM AZT, washed and pre-incubated with OCLN-derived peptides for 1 h. Monocytes were added at the top of a transwell with a confluent hCMEC/D3 monolayer and transmigration was allowed to happen for ≈ 16 h. Then, HIV RNA levels were measured in the upper and lower chambers of the transwell. (**E**) Schematic of the experimental setup. (**F**) Ratio of the amount of HIV RNA in the lower chamber was divided by the amount measured in the upper chamber. Each dot represents the ratio of the HIV RNA measurement performed in duplicate from a transwell containing a single organoid. The bar graph shows the mean +/- SEM. Two-tailed unpaired Student t test shows p value < 0.05 (*) or < 0.01 (**).

Together, our data indicates that the use of OCLN-derived peptides may represent a promising approach to decrease HIV neuroinvasion, and could help to decrease the burden of the CNS viral reservoir in seropositive patients.

## Discussion

The migration of monocytes from the bloodstream toward tissues – i.e. transmigration – is a tightly regulated event that requires the coordinated action of numerous proteins to allow the attachment of the monocyte to the endothelial wall, rolling and firm adhesion, interplay with endothelial cells to create a narrow “tunnel” across this physical barrier (either paracellular or transcellular), dramatic cell reshaping to squeeze through it, and release on the other side. Our study highlighted that monocytic OCLN favors monocyte transmigration, probably through its interaction with endothelial OCLN during the transmigration process. We identified OCLN-derived peptides that inhibit human monocyte transmigration *in vitro* and in zebrafish, and prevented monocyte-mediated HIV-1 neuroinvasion.

Our initial siRNA mini-screen showed that the absence of mOCLN decreased monocyte transmigration. It was surprising to observe that OCLN, a transmembrane protein known to promote tight cell-cell contact and a major element of TJs, in monocytes. However, we confirmed the present of OCLN in THP-1 cells and human primary monocytes by western blot. Another group detected OCLN protein in monocytes, reporting that the protein was overexpressed upon exposure to human cytomegalovirus (30). This observation was associated to increased infected monocyte migration, although they did not provide experimental evidence for a link between OCLN and migration at the time. OCLN has been shown to regulate the directional migration of epithelial cells (31). In this work, the authors indicate that OCLN would act as an actin scaffold protein at the leading edge of migrating cells. Although this discovery was made in adherent epithelial cells, which are very different from circulating monocytes, it is possible that OCLN also participate in the reorganization of the actin cytoskeleton during monocyte transmigration.

Mechanistically, several adhesion molecules (integrins, selectins, immunoglobulin-like superfamily) and chemokine receptors have been involved in monocyte transmigration (3). However, subcellular insights of the transmigration process are relatively scarce. A seminal study from 1998 investigated the distribution of F-actin and β-catenin in transmigrating rat monocytes (32), and the advent of intravital imaging recently allowed further understanding of this dynamic process (33). However, this later strategy deals with non-human monocytes, has low spatiotemporal resolution and could not readily provide access to the molecular mechanistic associated to this event. Our imaging analyses revealed that mOCLN can form a ring at monocyte-endothelial contact sites. Such imaging is very challenging and we could not reliably quantify and time this event, although we could reproducibly observe it.

Dendritic cells were shown to send dendrites outside the gut and airway epithelia, while expressing OCLN (34, 35), but no functional analyses were performed at the time. We showed here that silencing or overexpressing mOCLN impacts monocyte transmigration through endothelial cells. We hypothesize that mOCLN can form homotypic interactions with endothelial OCLN in order to squeeze through endothelial TJs while preserving endothelial permeability. Although this is an attractive working model, further investigations are required to determine whether these OCLN-mediated intercellular interactions are sufficient to locally retain full impermeability. Moreover, we observed mostly paracellular transmigration, i.e. migration of the monocyte between endothelial cells, but it remains unclear at this stage whether OCLN could also be involved in transcellular migration, a process during which monocytes would migrate through an endothelial cell (36).

We highlighted the transient appearance of an internal compartment in monocytes during the transmigration process. This compartment does not contain the classical markers of recycling (Rab11) nor endolysosomal compartments (Rab7). Moreover, the small GTPase Rab13 was previously shown to recycle OCLN (37), but we failed to show any colocalization between Rab13 and OCLN in human monocytes, neither at steady-state or during transmigration. We hypothesized that this compartment could actually represent a deep invagination of the plasma membrane, but fluid-phase marker could not reach this OCLN-containing compartment and thus, it likely represents a bona fide intracellular vesicle. Further investigations are ongoing to determine the nature and function of this compartment during monocyte transmigration.

We designed peptides that mimic the EL1 or EL2 of OCLN. Previous work showed that two peptides derived from xenopus OCLN altered epithelial impermeability (22). Another study showed that a peptide targeting the EL2 of rat OCLN but not EL1, perturbed the endothelial blood-testis barrier in a reversible manner (23). Here, our data shows that our newly designed peptides had no significant effects on vascular permeability *in vitro* and in zebrafish embryos. This difference could be attributed either to the difference of the peptide sequences, the different cell types investigated, and/or the different species used. Antibodies targeting the extracellular domain of OCLN have been developed (38), as they represent an interesting strategy to block Hepatitis C virus (HCV) entry in hepatocytes. Indeed OCLN is an essential entry factor for HCV (39), which drives the dynamics of viral particle internalization (40). Unfortunately, we did not manage to get access to these proprietary antibodies targeting the Els of OCLN. Nevertheless, the EL1 and EL2 peptides presented here could be tested as HCV antivirals.

We showed the ability of EL2 to inhibit human primary monocyte transmigration in zebrafish. We previously demonstrated the suitability of this xenotypic human-zebrafish model to study monocyte transmigration in the context of Zika virus infection (25), and another group also showed the survival and differentiation of human monocytes in zebrafish embryos (41). Here, we chose to use the human OCLN-derived EL1 and EL2 peptides together with human monocyte injection, which showed promising anti-transmigratory effects in vivo. More potent activity might have been seen if the homology between the zOCLN and human OCLN was greater, but as a proof-of-concept, our data highlights for the first time that targeting OCLN *in vivo* can represent an attractive approach to transiently modulate monocyte infiltration. Further immunological studies should provide more information as of whether an EL2 peptide treatment could attenuate local tissue inflammation.

HIV-associated neurocognitive disorders (HAND) are probably among the best characterized neuropathological symptoms described for a virus (5). In recent years however, evidences pointed toward macrophages as the main HIV reservoir, even under ART (42–46). The importance of the myeloid lineage as a master reservoir has also been highlighted by the fact that only patients transplanted with CCR5 Δ32/Δ32 haematopoietic stem-cells non-permissive to “macrophage-tropic” virus were were cured from HIV-1, while patients transplanted with wildtype CCR5 cells experienced viral rebounds after ART interruption (47–49)). The initial establishment of macrophages as HIV reservoirs in the CNS and other tissues is not a well-understood process. It is supposed that HIV takes advantage of blood-circulating monocytes that cross the vascular endothelium and enhances monocyte transmigration using a “Trojan horse” strategy (6). Here, we propose an innovative strategy, complementary to ART treatment, not aiming at targeting the virus, but rather at preventing it from reaching its brain reservoir. Indeed, we showed that EL1 or EL2 peptide treatments of transmigrating monocytes exposed to HIV resulted in more than ten time less HIV RNA detectable in cerebral organoids. Interestingly, in this experiment, the EL1 peptide also exhibited a strong effect, suggesting that the OCLN-derived peptides act on various steps to inhibit HIV neuroinvasion. Another team previously found that HIV-1 infection of monocytes is associated to increase expression of the TJAP molecules JAM-A and ALCAM (50). We were not able to see an increase of OCLN expression upon HIV-1 infection, but because mOCLN acts as a general transmigration factor, rather than a virus-specific one, one could anticipate that targeting mOCLN would have an impact on HIV-exposed monocyte transmigration.

Finally, we showed that treatment with AZT did not affect the outcome of HIV neuroinvasion, a particularly relevant finding. Indeed, monocytes are rarely productively infected in patients, but they can carry high amounts of infectious viral particles in intracellular vesicles. These protected virions would hence be indifferent to AZT, and remain infectious upon ART clearance. As the penetration of ART in the CNS is difficult, preventing HIV from invading the brain through the targeting of mOCLN represents an innovative strategy to reduce the HIV reservoir and/or HANDs in patients, which should be further explored.

## Material & Methods

### DNA constructs

An “all-in-one” Lentivector coding for two gRNAs targeting the OCLN gene, spCas9, E2-Crimson far-red fluorescent protein, and puromycin resistance was used. The initial plasmid backbone used was the pLenti-spCas9-T2A-Crimson-P2A-puro (kindly provided by Dr. Feng Zheng) and derived versions of this plasmid were also generated to express mNeonGreen (Addgene #122183), TagRFP-T (Addgene #122200) or iRFP670 (Addgene # 122182) fluorescent proteins. Here, the pLenti-spCas9-T2A-Crimson-P2A-puro was cleaved using the BsmBI restriction enzyme (Thermofisher) and treated with FastAP (Thermofisher). A synthetic DNA coding for BsmB1 enzyme, a first gRNA targeting the OCLN gene (5’-GGCCTCTTGAAAGTCCACCT-3’) under the control of the human U6 promoter, a tracrRNA for Cas9 recognition, a termination sequence, a second gRNA targeting the OCLN gene (5’-TGTCATCCAGGCCTCTTGAA-3’) under the control of the hH1 promoter, a tracrRNA, and a BsmBI sequence was synthesized (Integrated DNA Technologies). The fragment was amplified using Phusion polymerase (Thermofisher). The insert was cloned into the prepared backbone using the T4 DNA fast ligase kit (Thermofisher) and the product was transformed in Stbl3 bacteria. The final lentivector pLenti-gRNA-OCLNx2-spCas9-T2A-Crimson-P2A-puro used in this study and depicted in Figure 1E is available on Addgene (#208398) as well as the variants pLenti-gRNA-OCLNx2-spCas9-T2A-mNeonGreen-P2A-puro Addgene (#208400) and pLenti-gRNA-OCLNx2-spCas9-T2A-iRFP670-P2A-puro Addgene (#208399).

### Antibodies

List of the used antibodies, types of experiments where they were used and their dilutions:

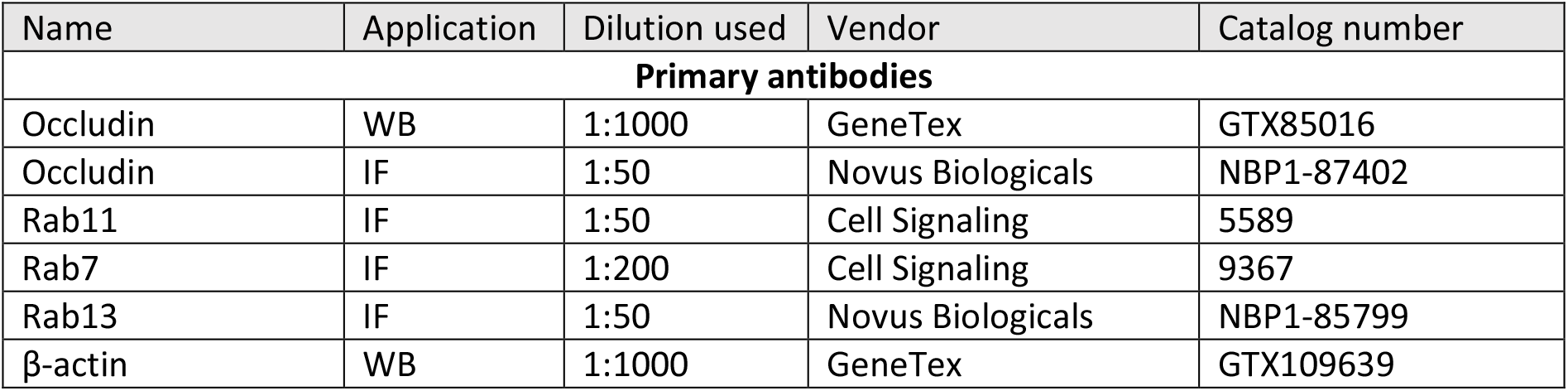
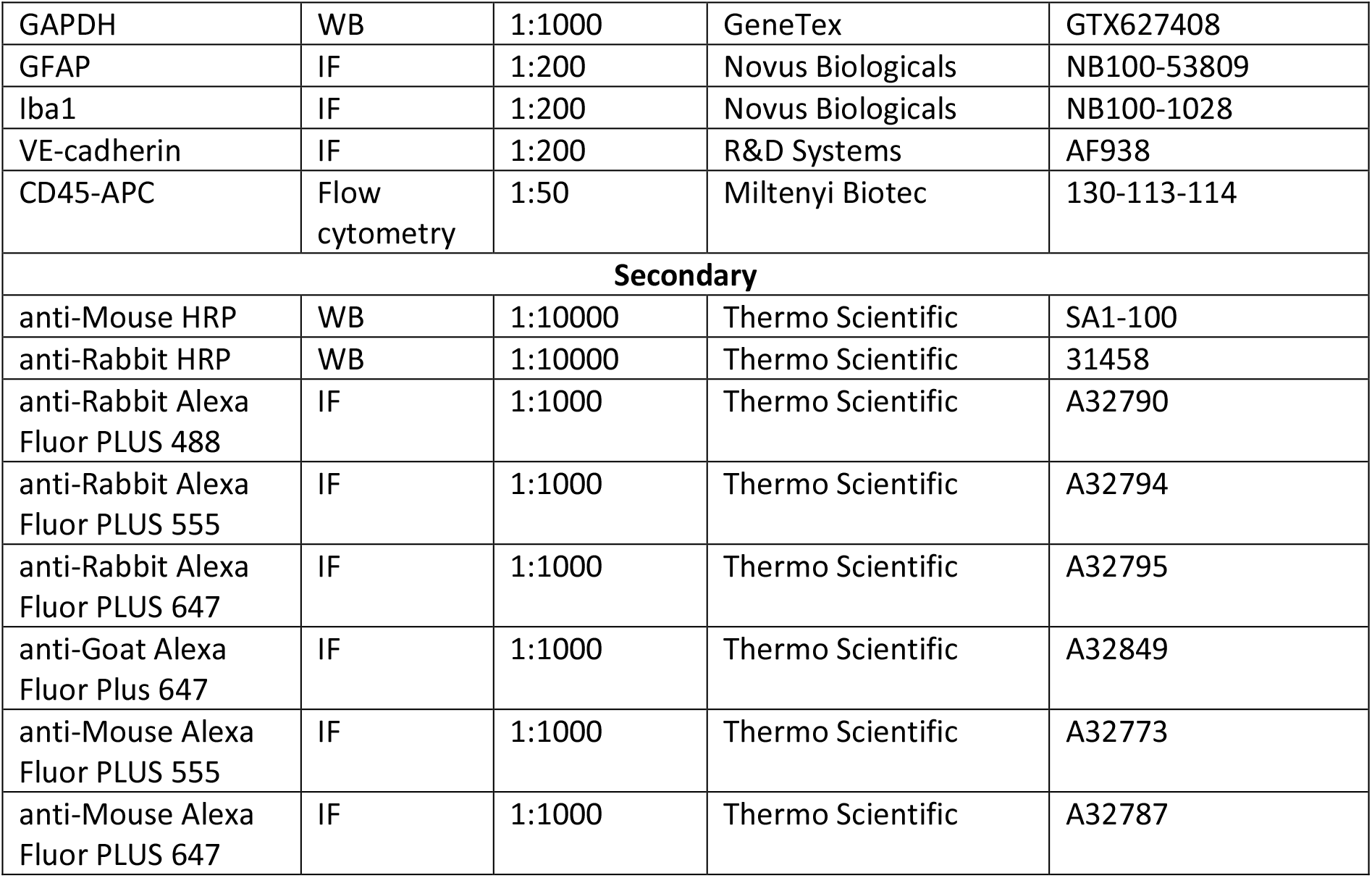

### Cell culture

Human primary monocytes and THP-1 cell lines (ATCC) were cultured in RPMI 1640 (Gibco) supplemented with 10% FBS (Sigma) and 1% Penicillin-Streptomycin (PS; Gibco). Human cerebellar microvascular endothelial cells D3 (hCMEC/D3) were kindly provided by Dr. Sandrine Bourdoulous (Institut Cochin, Paris, France). They were grown in EndoGro-MV Complete culture medium (Millipore) supplemented with 1 ng/mL b-FGF (Sigma) and 1% Penicillin-Streptomycin (Gibco) on 200 μg/mL rat collagen type 1-coated (Sigma) plates. For experimental use of hCMEC/D3 cells, EndoGro-MV complete medium was additionally supplemented with 10 μM Resveratrol (Sigma) and 10 mM Lithium Chloride (Millipore); cells were differentiated for 7 days prior to an experiment. HEK 293T cells were cultured in DMEM (Gibco) supplemented with 10% FBS and 1% PS. Mouse fibroblast cells (MEF) feeder cells were kept in DMEM with 10% FBS, 1% PS and 1 mM Sodium Pyruvate (Sigma). All cells were kept at 37°C in 5% CO_2_.

### Human monocyte isolation

Peripheral blood mononuclear cells (PBMCs) were isolated from buffy coats of healthy blood donors provided by EFS Montpellier, France. All donors signed a consent form allowing the use of their blood for research purposes. PBMCs were purified on a density gradient with Ficoll-Paque (Cytiva). Monocytes were isolated with CD14^+^ microbeads and LS columns (Miltenyi Biotec). Monocytes from a total of 32 donors were used.

### Transcriptomic analysis of monocytic TJAPs

RNA-seq data from primary human monocytes and THP-1 cells were downloaded from the Gene Expression Omnibus (GEO) (15). RNA-seq read counts from four monocyte donors (accession ID: GSE74246) were further normalized to reads per kilobase of exon model per million mapped reads (RPKM) in R. The RNA-seq data from THP-1 cells (accession ID: GSM927668) was already available in RPKM. The heatmap was created in Excel.

### SiRNA-based THP-1 transmigration screen

Monocytic TJAPs were knocked down (KD) in THP-1 cells via electroporation using a siGENOME siRNA pool (RNAi Cherry-picked Library, Dharmacon LP_29911). A non-targeting siRNA (siGENOME Non-Targeting siRNA Pool #2, Dharmacon D-001206-15-05) was used as a control (siCtrl). THP-1 cells were counted, centrifuged (1000 rpm, 5 min, RT), and suspended in cold PBS to a cell density of 5×10^6^ cells/ml. The cells were then aliquoted in Eppendorf tubes at a concentration of 10^6^ cells/tube in 200 μL. The siRNA was added to a final concentration of 200 nM. The cells were gently mixed with the siRNA and incubated for 5 min on ice. The cell/siRNA mixture was transferred to an electroporation cuvette (2 mm). The cells were electroporated at RT with the following parameters: 600 V, 50 µs, 4 pulses, interval 1 s using an ECM 830 Square Wave Electroporation System (BTX No. 45-0052). The electroporated cells were taken out of the cuvette using a sterile plastic Pasteur pipette, placed in Eppendof tubes, and incubated for 10 min on ice. Finally, the cells were seeded in 12-wells plates containing cell culture media without antibiotics (1.5 ml/well of RPMI 10% FBS, 1 mM Sodium pyruvate). Non-electroporated cells were used as negative control.

The transmigration experiment was performed in 96-wells transwells (Corning, 5 μm pore size, PC). The hCMEC/D3 cells were grown as described above. The KD THP-1 cells were harvested after 3 days of electroporation. A total of 150 000 cells were added per transwell in a volume of 100 μL of cell culture media (RPMI, 10% FBS, 1 mM sodium pyruvate, 1% PS), while freshly prepared media (RPMI, 10% FBS, 1% PS) supplemented with 200 ng/ml MCP-1 (R&D Systems) was added to the bottom chamber. After an overnight incubation (approximately 17 h), the transmigrated cells present in the bottom chamber were harvested and counted. The percentage of transmigrated cells was calculated with respect to the siCtrl condition.

### Lentivirus production

All viruses and lentiviruses were produced by transfecting Hek293T cells using CalPhos Mammalian Transfection Kit (Takara Bio) according to the manual in DMEM, 10% FBS, 1% PS. For viral production, a single plasmid was used (100-200 ng/cm^2^). For lentiviral production, the ratio of 2:1:2 was used for construct-of-interest: VSVG:psPAX2 (200-300 ng/cm^2^ total). Overnight after transfection, medium was changed to fresh DMEM, 10% FBS, 1% PS. Medium containing viruses or lentiviruses was harvested 48 h later. It was spun down (1000 g, 5 min) and filtered through a syringe filter with 0.45 μm pores. Then, the supernatant was transferred onto 10% sucrose (Sigma) in PBS inside a 32 mL open-top thickwall polycarbonate ultracentrifuge tube (Beckman Coulter) and centrifuged using SW32 Ti rotor in Optima XE-90 ultracentrifuge (Beckman Coulter) at 28000 rpm for 2 h at +4°C. The supernatant was discarded and pellets were resuspended in cold RPMI (1/100) on ice. After 1 h of incubation on ice, RPMI with viruses/lentiviruses was frozen at -80°C until used. The titer of lentiviruses was assessed by flow cytometry using the fluorescent tag of the protein of interest.

### HIV production and titration

Plasmids NLAD8 HIV-1 AD8 Macrophage-Tropic R5 (NIH AIDS reagents #11346) or HIV Gag-imCherry V3 loop (29) were transfected in HEK 293T cells using CalPhos Mammalian Transfection Kit (Takara Bio) in DMEM, 10% FBS, 1% PS, according to the manufacturer’s instructions. After overnight incubation, the media was changed to fresh media. The supernatant containing the viruses was harvested 72 h later. It was spun down (1000 g, 5 min) and filtered through a syringe filter with 0.45 μm pores. Then, supernatant was transferred onto 10% sucrose (Sigma) in PBS in a 38.5 ml thin-wall ultra-clear ultracentrifuge tube (Beckman Coulter) and ultracentrifuged using SW32 Ti rotor in Optima XE-90 ultracentrifuge (Beckman Coulter) at 28 000 rpm for 2 h at 4°C. Supernatant was discarded and pellets were resuspended in cold RPMI on ice for 4 h. Virus stocks were frozen at -80°C until used. The titer of viruses was assessed at 48 hpi by flow cytometry on GHOST X4/R5 cells (NIH AIDS Reagents # 3942; (51)).

### Peptides

OCLN-derived peptides were designed to represent a part of each extracellular loop (EL) of OCLN. EL1-derived peptide: DRGYGTSLLGGSVGYPYGGSGFGS and EL2-derived peptide: YGSQIYALCNQFYTPAATGLYVD. They were generated based on a study from Tavelin et al (21). They were designed to cover the potential site of Occludin-Occludin interaction. Scramble peptides were generated by randomly reordering the amino acids of the peptides. The scrEL1 used in this study was: GGTSPLYGGFVGGYSGDSYGRGSL and the scrEL2 peptide was: PQAYDFTGNGSYLCTLYAYVIAQ. The peptides were synthesized by JPT Peptide Technologies and ProteoGenix.

### Transmigration assay

For transmigration experiments which do not involve HIV, hCMEC/D3 cells were seeded in cellQART 24-well transwell inserts (Sabeu, 5 μm pore size PET) at 30 000 cells/insert and grown for 7 days on collagen as described above. Primary monocytes or THP-1 cells were added at the top of the transwell (200 000 cells/well, 0.2 ml of RPMI, 10% FBS, 1% PS) while freshly prepared media (RPMI, 10%, FBS 1% PS) supplemented with 200 ng/ml MCP-1 (R&D Systems) was added to the bottom chamber. Monocytes or THP-1 cells were transduced 48 h before the experiment, washed in RPMI 10% FBS 1% PS once after overnight incubation and once before the transmigration experiment. In the case of OCLN-derived peptide, monocytes were washed in RPMI 10% FBS 1% PS once after overnight incubation, incubated with OCLN-derived peptides and the monocyte-peptide mixture was used the next day for the transmigration experiment. The transmigration through hCMEC/D3 cells was performed overnight, and cells from the bottom and the top chambers were recovered and prepared for flow cytometry analysis.

For transmigration experiments which involve HIV, hCMEC/D3 cells were seeded in 96-well transwells (Corning, 5 μm pore size, PC) at 15 000 cells/insert and grown for 7 days on collagen as previously described. After 7 days, hCMEC/D3 cells were washed, and monocytes were added at the top of the transwell after 1 h incubation with OCLN-derived peptides (100 000 cells/well, 0.1 mL of RPMI, 10% FBS, 1% PS). The bottom chamber was filled either with 0.22 ml of fresh media (RPMI, 10% FBS, 1% PS) supplemented with 200 ng/mL of MCP-1 or with a 35+ days-old brain organoid in 0.22 ml of Neurobasal medium with N2 supplement, 1x GlutaMAX, 1% PS. After 24 h of incubation the top and bottom chambers were lysed for RNA extraction and RT-qPCR.

### Adhesion assay

Adhesion assay was performed in a white cell-culture treated 96-well plate with a flat bottom (Corning). The coating of the bottom of the plate was done with 150 μg/ml of collagen type I solution from rat tail, or 10 μg/mL of human fibronectin. Monocytes were added to the coated substrate and incubated for 1 h at 37 °C. Then, monocytes were washed 2x with PBS to remove non-adherent cells. The cells that remained attached were analyzed with the CellTiter-Glo Luminescent Cell Viability Assay (Promega).

### Immunofluorescence and image acquisition

THP-1 cells or primary human monocytes were seeded on to 12 mm glass coverslips in 24-well plates, with either a 10 μg/ml fibronectin (R&D Systems) coating or a confluent monolayer of adherent hCMEC/D3 cells, and fixed in 4% PFA for 15 min at room temperature (RT). Cells were permeabilized with 0.05% triton X-100 (Sigma) with 0.5% BSA (Euromedex) for 15 min, then blocked in 10% FBS (Sigma) with 1:5 human FcR blocking reagent (Miltenyi) for 20 min. Primary antibody staining was performed for 1 h in PBS (Gibco), followed by washes and secondary antibody staining for 1 h. Images were acquired using an Andor Dragonfly Spinning Disk Microscope (Oxford Instruments) equipped with a 1024×1024 EMCCD camera (iXon Life 888, Andor) with either a 40X, 63X or 100X objectives, or a Confocal LSM 980 (Zeiss) with a 63X objective. Pre-processing of images was performed in either FUSION (Oxford instruments) for Dragonfly or ZEN Blue for the LSM 980 microscopes.

The hESC-derived cerebral organoids were fixed in 4% PFA overnight at 4°C and permeabilized with 0.5% triton X-100 with 0.5% BSA for 24 h. Blocking was done with 0.5% BSA supplemented with 1:5 human FcR blocking reagent for 4 h followed by 24 h staining firstly with primary antibodies and then secondaries in 0.5% triton X-100 with 0.5% BSA. Organoids were transferred to a black 96-well µ-plate (ibidi) and cleared in RapiClear 1.52 reagent (Sunjin Lab) to perform deep fluorescence imaging. Images were acquired using an Andor Dragonfly Spinning Disk Microscope.

### Live-cell imaging

For transmigration experiments, hCMEC/D3 cells were grown for 7 days in their experimental medium on collagen as described above in a 35 mm imaging dish with 4 compartments with a polymer coverslip bottom (Ibidi), then stained with CellTrace Yellow (Invitrogen) 1 h before the beginning of imaging for 20 min and washed 3 times with RPMI 10% FBS 1% PS. THP-1 cells or primary human monocytes previously transduced with GFP+-constructs were put on pre-stained hCMEC/D3 cells 30 min before the beginning of imaging. For fluid phase marker imaging, fluorescent Dextran was used as previously (52). Briefly, 100 µg/ml 3kDa Dextran was added to… Imaging was done using Andor Dragonfly Spinning Disk Microscope as above with 100x objective (frame rate: 10 min, z step: 0.8 µm).

### Image processing and quantification

Image processing was performed in Imaris software v9.7 (Bitplane, Oxford Instruments) or the Fiji upgrade of ImageJ. Co-localization was assessed using Imaris software using standard procedure. Briefly, images were threshold using EGFP fluorescence intensity to remove background and diffuse signal. Threshold was kept the same per condition and experiment repeats. Pearson correlation coefficient was and plotted for comparative analyses.

### Flow cytometry

Cells expressing fluorescent proteins were fixed in 4% PFA and washed in PBS before flow cytometry. When immunostaining was used, fixed samples were blocked with 10% FBS in PBS and 1:5 FcR Blocking Reagent. To label cell surface proteins, samples were stained with primary conjugated antibodies on ice for 1 h and washed 2x with PBS. Samples were acquired with NovoCyte Flow Cytometry System (ACEA Biosciences). Data was analyzed using FlowJo (LLC).

### Western Blotting

Cells were lysed in RIPA buffer at about 10^6^ cells in 50 µL (50 mM Tris-HCl (Sigma) pH 8.0, 150 mM NaCl (Honeywell), 1% NP40 (Sigma), 0.5% sodium deoxycholate (Sigma), 0.1 % SDS (Sigma), freshly added protease inhibitor cocktail 1x (Promega)). Samples were sonicated at 4°C in a waterbath ultrasonic cleaner (Velleman) for 8 min, denaturated in NuPage LDS Sample buffer (Invitrogen) supplemented with DTT (Thermofisher) at final concentration 100 mM for 3 min at 98°C. Samples were run on Bolt 4-12% Bis-Tris Plus Gels (Thermofisher) at 140V for 1.5 h in MES Running Buffer (Thermofisher), then transferred onto Trans-Blot Turbo Mini PVDF membrane 0.2 μm (Bio-Rad) using Trans-Blot Turbo Transfer System (Bio-Rad). Samples were blocked in 5% milk (Regilait) in TBS supplemented with 0.05% Tween 20 (Sigma; TBST). Membranes were incubated with primary antibodies at 4°C overnight in TBST 0.05% with 5% BSA (Euromedex), washed 3x 10 min with TBST 0.05%, incubated with secondary HRP antibodies for 1 h at RT, washed 3x 10 min with TBST 0.05% and revealed using either Clarity or Clarity Max ECL Substrate (Bio-Rad). Chemiluminescence imaging was performed using a ChemiDoc Imaging System (Bio-Rad).

### RT-qPCR

RNA was extracted either from cells and organoids using the RNeasy Kit (Qiagen) or from supernatants using NucleoSpin RNA virus kit (Macherey-Nagel). Reverse transcription and quantitative real-time PCR was performed using Luna Universal One-Step RT-qPCR Kit (New England Biolabs) in a 96-well plate and qPCR was performed in a LightCycler 96 System (Roche). Data was not normalized to β-actin because only HIV relative quantity was needed. β-actin was used to assess the presence of cells. HIV gag primers: 5’-ACTCTAAGAGCCGAGCAAGCT-3’, 5’-TCTAGTGTCGCTCCTGGTCC-3’. β-actin primers: 5’-CACCATTGGCAATGAGCGGTTC-3’, 5’-AGGTCTTTGCGGATGTCCACGT-3’.

### Lucifer yellow permeability assay

hCMEC/D3 cells were cultured as described before for 7 days in transwells before any experimental manipulations. For monitoring hCMEC/D3 permeability with Occludin-derived peptides, peptides were added overnight and not washed away before Lucifer yellow addition. Permeability was assessed by adding 50 μM Lucifer yellow in the top chamber. After 2 h incubation inside the incubator, media was collected from the top and bottom chambers of transwells. Quantification of fluorescence was done using a microplate reader (Infinite 200 Pro M Plex, Tecan) with excitation at 428 nm and emission at 536 nm.

### Zebrafish embryo permeability and transmigration assay

Tg(fli1a:eGFP-CAAX) zebrafish (*Danio rerio*) embryos were used in all experiments. Embryos were maintained in Danieau 0.3X medium (17.4 mM NaCl, 0.2 mM KCl, 0.1 mM MgSO_4_, and 0.2 mM Ca(NO_3_)_2_) buffered with 0.15 mM HEPES (pH = 7.6) and supplemented with 200 mM of 1-Phenyl-2-thiourea (Sigma-Aldrich) to inhibit the melanogenesis. For all zebrafish experiments, the offspring was selected based on anatomical/developmental good health. Embryos were split randomly between experimental groups. Forty-eight hours post fertilization (hpf), embryos were mounted in a 0.8% low-melting-point agarose pad containing 650 mM of tricain (ethyl-3-aminobenzoate-methanesulfonate) to immobilize them. Primary human monocytes frozen at 10.10^6^ cells/mL were thawed 2h and seeded at 5.10^6^ cells/mL before their injection in zebrafish. Monocytes were labeled with CellTrace Calcein Red-Orange (Invitrogen) according to the manufacturer’s instructions. Monocytes were resuspended in PBS in which the peptides EL1 or EL2 or DMSO were diluted at 1:100. Monocytes were injected with a Nanoject microinjector 2 (Drummond) and microforged glass capillaries (20-µm inner diameter) filled with mineral oil (Sigma-Aldrich), and 18.4 nL of a cell suspension at 10^8^ cells/mL were injected in the duct of Cuvier of the embryos under a M205 FA stereomicroscope (Leica). Injected embryos were maintained at 28°C in between injection and imaging. At 6–8h post injection, confocal imaging was performed with an inverted Olympus Spinning Disk (30X objective, NA 1.05). Image analysis and processing were performed using ImageJ.

### Stem cell culture and cortical organoid differentiation

H9 human embryonic stem cells (hESCs; female, WA09, WiCell) were cultured in mTeSR Plus medium (STEMCELL Technologies) with 1% Penicillin-Streptomycin in 8 mg/mL Matrigel-coated (Corning) 6 cm dishes (Corning). The medium was changed daily and cells were split by cutting colonies every 5-6 days. For hESC-derived cerebral organoid differentiation hESCs were transferred while splitting on MEF feeder layer in ESC medium (DMEM F12 + L-glutamine (Gibco), 20% KnockOut Serum Replacement (Gibco), 1x MEM Non-Essential Amino Acid Solution (Gibco), 100 μM β-mercaptoethanol (Gibco), 1% PS) with freshly added 8 ng/mL human bFGF (Sigma). The medium was changed daily. When hESC colonies reached 1.5-2 mm, medium was replaced with differentiation medium (ESC medium with 1 mM Sodium Pyruvate, no bFGF), then they were picked and lifted using a cell lifter (Corning) and transferred to a 6 cm ultra-low attachment dish (Corning) together with differentiation media. The medium was replaced every other day. On day 0 (24 h post-lifting, when embryoid bodies were formed) half of the medium was replaced with differentiation medium supplemented with 3 μM IWR-1-Endo (Sigma), 1 μM Dorsomorphin (Sigma), 10 μM SB-431542 (Sigma), and 1 μM Cyclopamine (Sigma) final concentration. This supplemented medium was maintained and changed every other day On day 3, differentiating organoids were placed on a tissue-culture orbital shaker in a tissue culture incubator. On day 18, medium was switched to Neurobasal medium (Gibco) with 1x N2 supplement (Gibco), 1x GlutaMAX (Gibco), 1% PS, and 1 μM Cyclopamine (Sigma), changed every other day. On day 24, Cyclopamine was removed from the content of the medium. Organoids were taken for experiments starting from day 35.

### Statistical analysis

Student’s t test was performed as described in figure legends. Analyses were done using GraphPad Prism (v9.5.1).

## Data and materials availability

All data are available in the main text, the supplementary materials or will be made available on reasonable request.

## Acknowledgments

We acknowledge the MRI imaging facility, part of Biocampus Montpellier, member of the national infrastructure France-BioImaging, for advice and training. The following reagent was obtained through the NIH HIV Reagent Program, Division of AIDS, NIAID: pNLAD8 HIV-1 AD8 Macrophage-Tropic R5 (11346); GHOST CXCR4+CCR5+ Cells (#ARP-3942), contributed by Dr. Vineet N. KewalRamani and Dr. Dan R. Littman. We thank Philippe Benaroch (Institut Curie, paris, France) from providing the HIV Gag-imCherry V3 loop DNA plasmid. We thank Isis Ricaño-Ponce (Innova Bio Data) for her help with the RNAseq data analyses and Lucie Klughertz for HIV-1-related experiments.

## Author contributions

Conceptualization: RG. Methodology: DB, NVAN, YB, EP, AD, VM, VL, JG, NO, RG. Investigation: DB, NVAN, YB, AD, NO. Analysis: DB, NVAN, YB, AD, NO, RG. Supervision: JG, NO, RG. Writing – original draft: RG. Writing – review & editing: DB, NVAN, YB, EP, AD, JG, NO, VL, RG.

## Supplemental Figures and legends

**Suppl. Figure S1.**
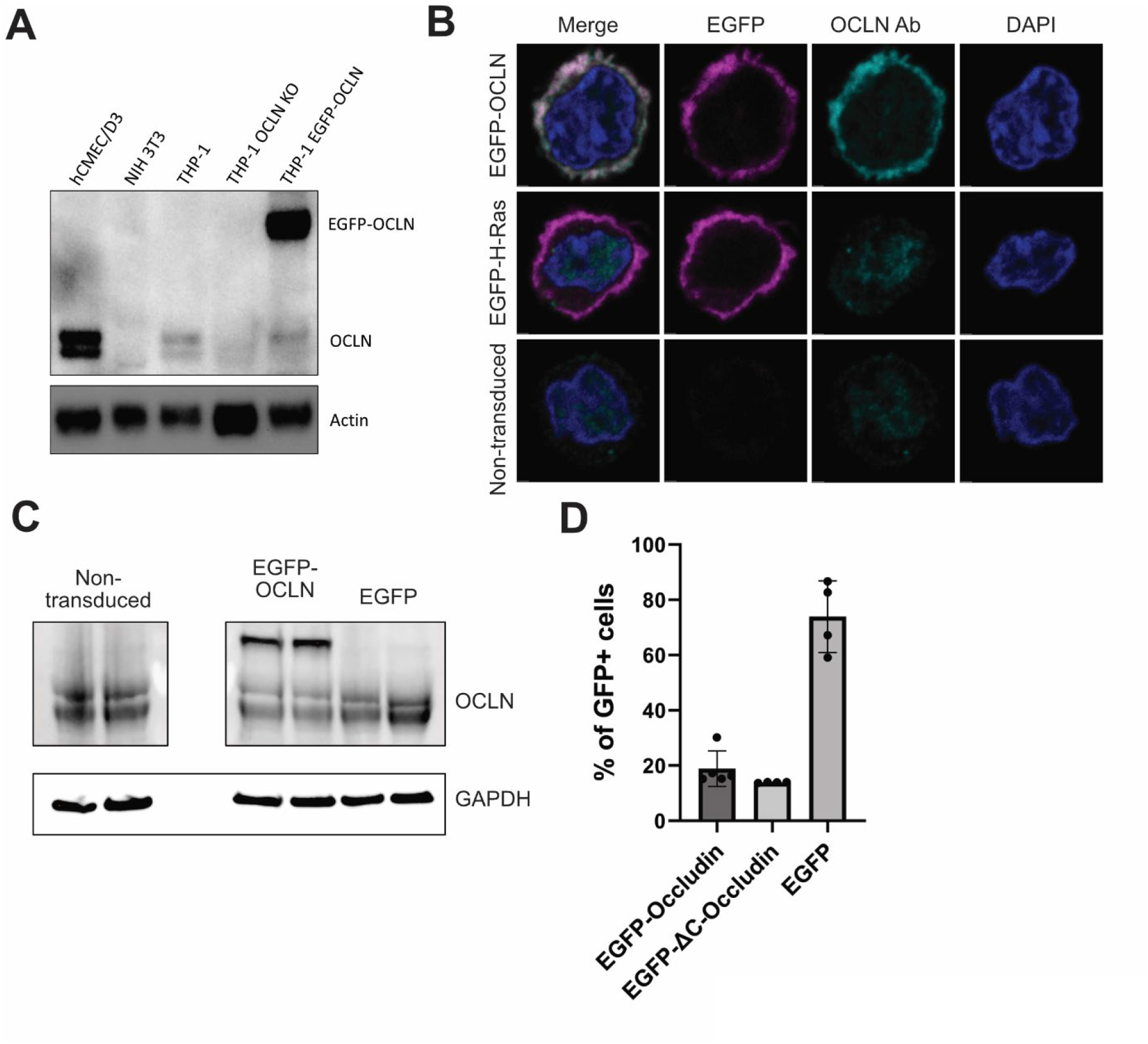
Characterization of monocytic OCLN expression. (**A**) Western Blot analysis of OCLN expression in indicated cell lines. Actin is used as a loading control. hCMEC/D3 and THP-1 cells express OCLN (two bands), the THP-1 OCLN KO cells do not show OCLN, although non-specific bands of lower size and very weak intensity can be observed. As controls, NIH 3T3 do not express OCLN and THP-1 transduced with EGFP-OCLN express both endogenous and overexpressed OCLN. (**B**) Immunofluorescence images of primary monocytes transduced with EGFP-OCLN, EGFP-CAAX, or non-transduced, fixed and stained with anti-OCLN antibody (cyan) and Dapi (yellow). EGFP is in magenta. Scale bar: 2 μm. (**C**) Western Blot showing endogenous OCLN expression by primary human monocytes and exogenous expression of EGFP-OCLN in the same donors. (**D**) Efficiency of transduction of primary monocytes from 2 independent experiments. Error bars are mean +/-SD.

**Suppl. Figure S2.**
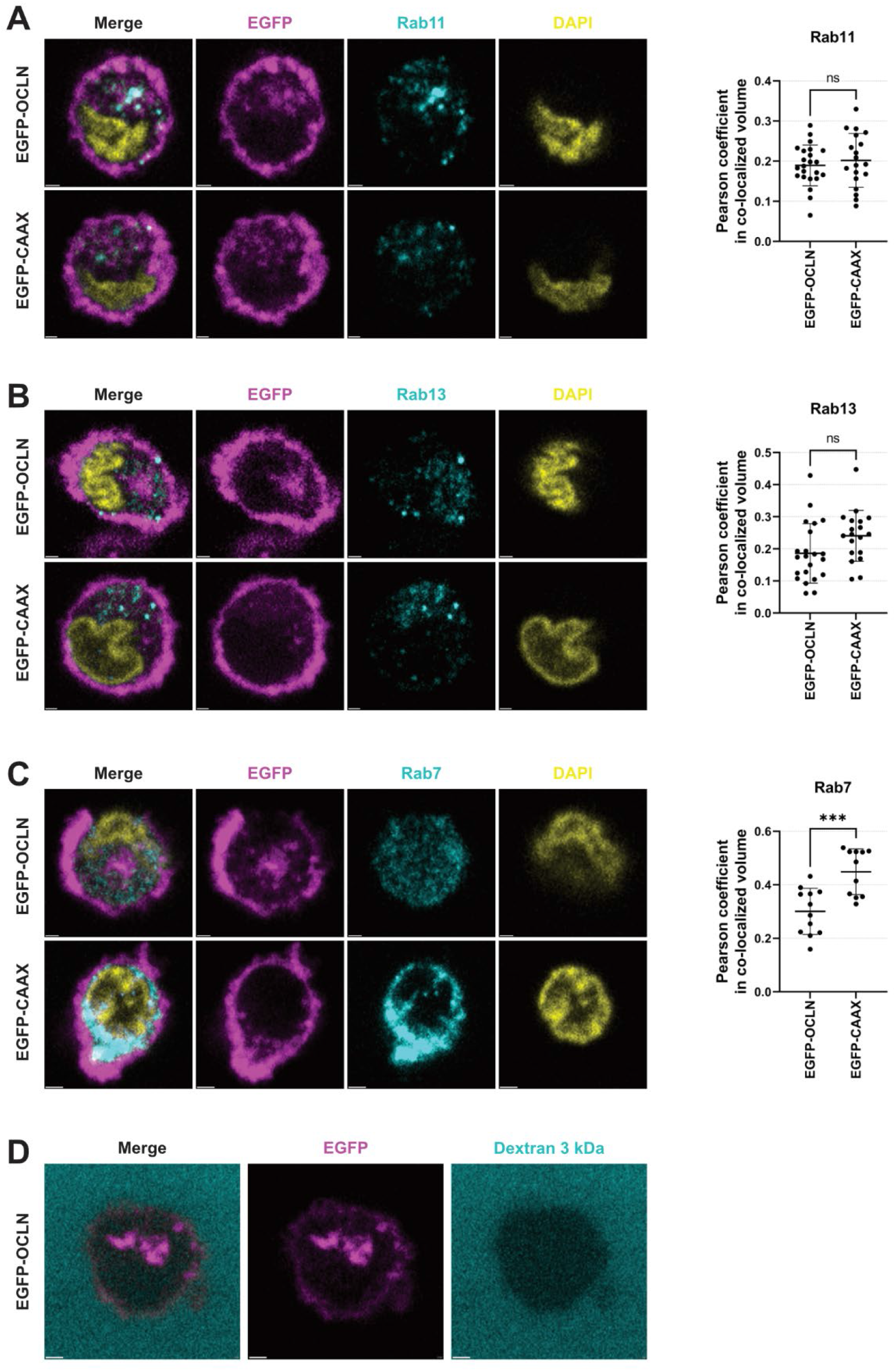
Characterization of the OCLN-containing compartment. (**A-C**) Immunofluorescence image of primary monocytes expressing either EGFP-OCLN or EGFP-CAAX (magenta) attached to hCMEC/D3 cells and stained with Dapi (blue) and antibodies (cyan) against Rab11 (**A**), Rab13 (**B**) or Rab7 (**C**). Scale bar: 1 μm, except in C for EGFP-CAAX: 2 μm. (**D**) Primary monocyte expressing EGFP-OCLN (magenta) attached to hCMEC/D3 monolayer were incubated with 3 kDa fluorescent Dextran (cyan) and immediately imaged using confocal microscopy. The snapshots highlight that Dextran does not access the OCLN-containing compartment. Scale bar: 1 μm.

**Suppl. Figure S3.**
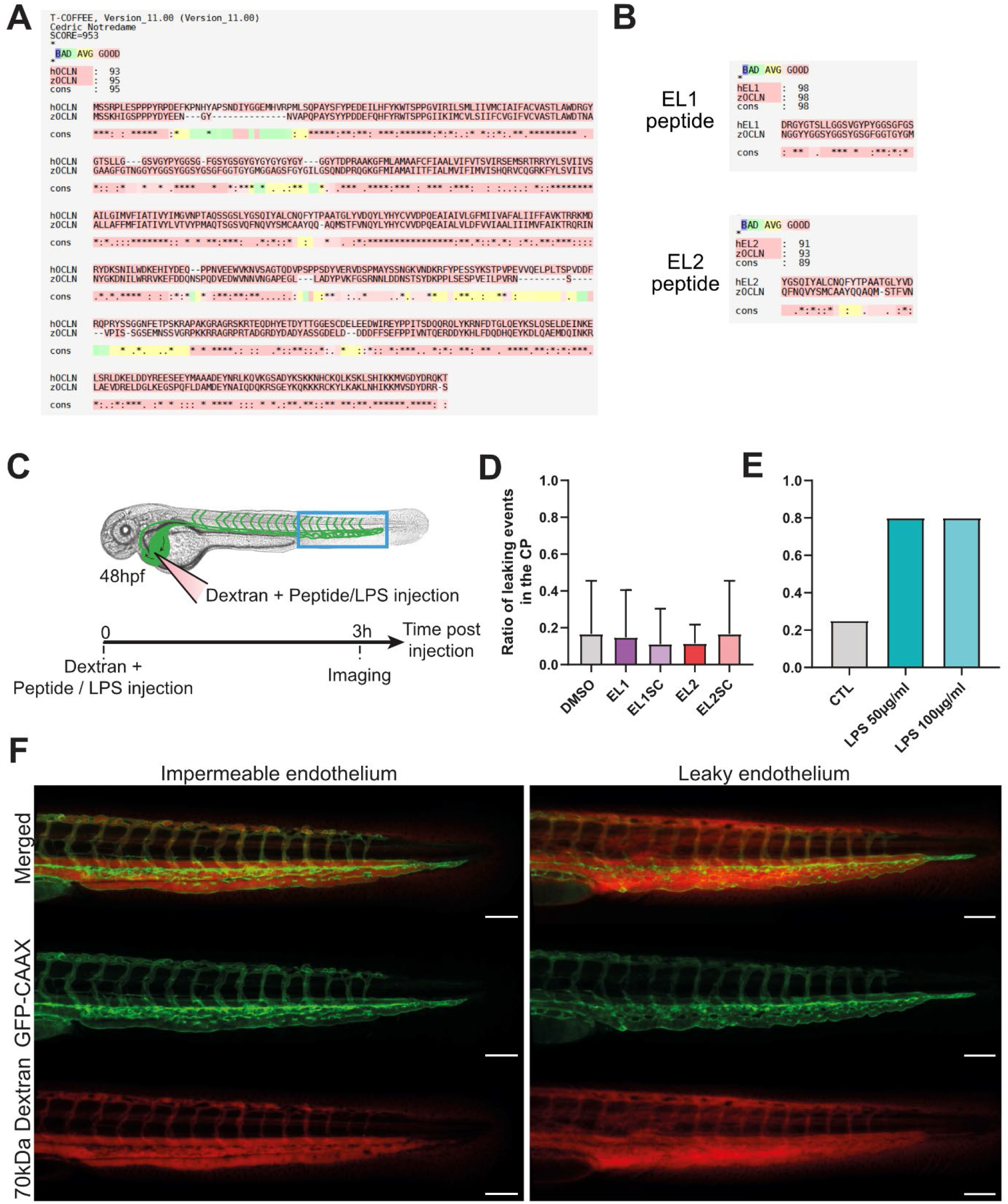
Characterization of the zebrafish embryo model. (**A**) Comparative analysis of the human OCLN (hOCLN) and *danio rerio* (zebrafish) OCLN (zOCLN) amino acid sequences and consensus sequence using T-Coffee (see Material & Methods for details). Red underlining of amino acids indicates good sequence similarity. (**B**) Comparative analysis of the EL1- and EL2-derived sequences of hOCLN with zOCLN. Red underlining of amino acids indicates good sequence similarity. (**C**) Schematic of the experimental procedure associated with the zebrafish model to test the effect of EL1, EL2 and their scramble peptides on endothelial permeability. Treatment with LPS is used as control for permeability inducing agents. (**D**) Scatter plot with bar of the ratio +/-SD of leaky endothelium at 3 h post dextran and peptide injection. The experiment was carried out three independent times. (n_DMSO_ = 39; n_EL1_ = 25; n_scEL1_= 26; n_EL2_ = 24; n_scEL2_= 23). (**E**) Scatter plot with bar of the ratio of leaky endothelium at 3 hours post dextran and peptide injection. The experiment was carried out one time. (n_Control_ = 12; n_LPS-50µg/mL_ = 15; n_LPS-100µg/mL_ = 15). (**F**) Representative images of zebrafish embryos with an impermeable or leaky endothelium in the tail at 3 hours post injection of PBS or LPS respectively. Scale bar: 100 µm.

**Suppl. Figure S4.**
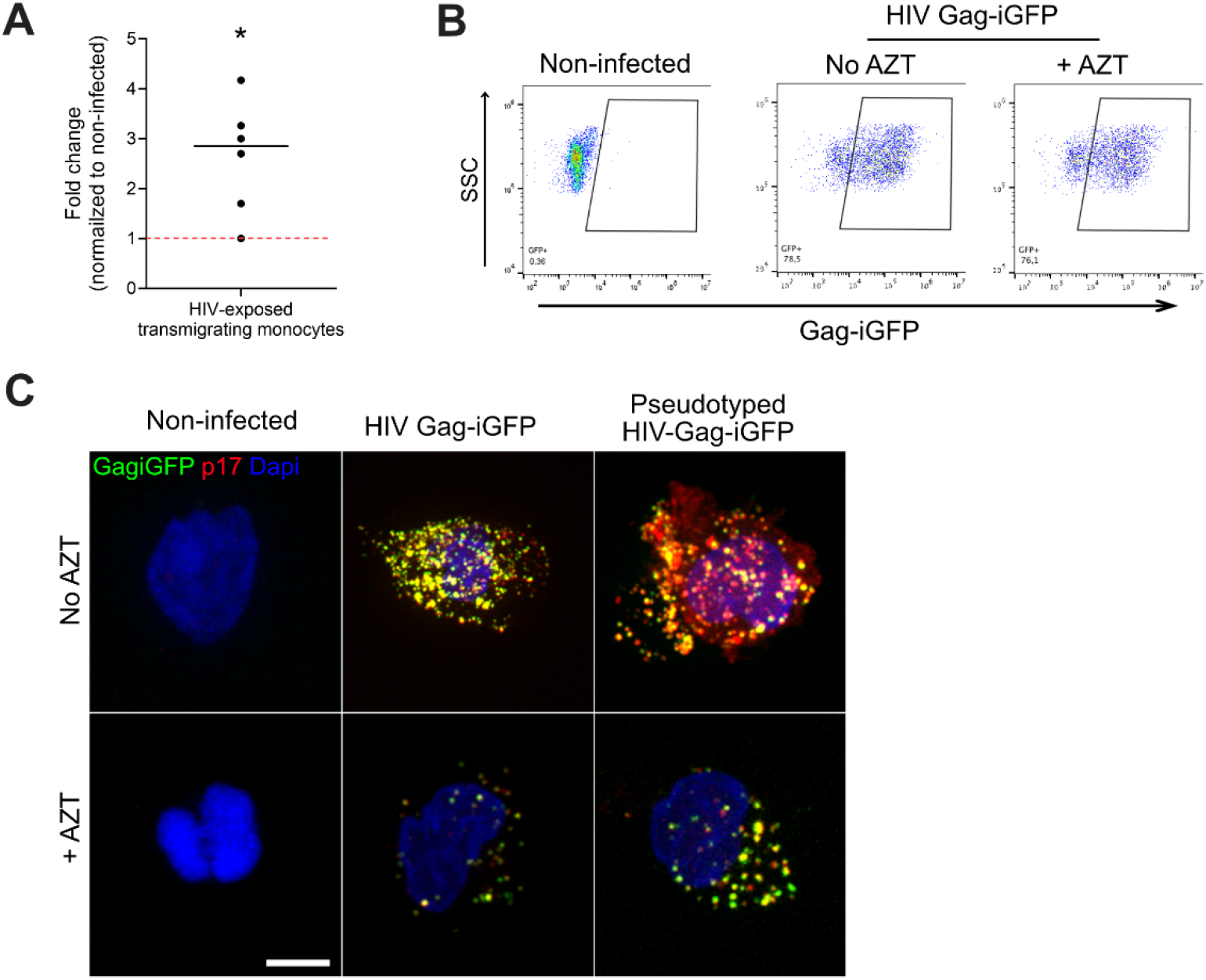
Characterization of primary monocyte infection by HIV-1 and transmigration. (**A**) Primary monocytes were infected for 48 h at MOI 1 with HIV-1 (NLAD8) and added to the top chamber of transwells containing a hCMEC/D3 monolayer to allow transmigration to occur. After overnight incubation, the bottom chamber was harvested and the number of transmigrated monocytes was counted. The dot plot shows the mean of the fold change of the number of HIV-exposed monocytes over non-infected counterparts. Each dot corresponds to a donor. Paired Student T test p value < 0.05 (*). (**B**) Primary monocytes were non-infected or infected with HIV-1 (NLAD8) at MOI 1 for 48 h in the presence or absence of AZT. The dot plots show the percentage of Gag-iGFP-expressing cells as a function of the side scatter (SSC) measurement as determined by flow cytometry. The data highlights that despite AZT treatment, monocytes are positive for Gag-iGFP as they carry fluorescent particles. (**C**) Primary monocytes were non-infected, infected with HIV-1 Gag-iGFP, or HIV-1 Gag-iGFP pseudotyped with VSV-G, at MOI 1 for 48 h in the presence or absence of AZT. Cells were fixed and stained for Gag p17 (red) and Dapi (blue) and Gag-iGFP is shown in green. Scale bar: 5 µm.

## Supplemental Movies

**Movie S1. Live cell imaging of an EGFP-OCLN transmigrating primary monocyte.** 3D time lapse spinning disk confocal imaging of a primary monocyte transduced with EGFP-OCLN (red) on hCMEC/D3 monolayer pre-stained with CellTrace (cyan) before the beginning of imaging. Z stacks images were taken every 10 min for 9 h and 30 min. Scale bar: 10 μm.

**Movie S2. Live cell imaging of an EGFP-OCLN transmigrating THP-1 cell.** 3D time lapse spinning disk confocal imaging of a THP-1 cell transduced with EGFP-OCLN (red) on hCMEC/D3 monolayer (not stained). Z stacks images were taken every 10 min for 9 h and 10 min. Scale bar: 10 μm.

**Movie S3. Live cell imaging of an EGFP-CAAX transmigrating primary monocyte.** 3D time lapse spinning disk confocal imaging of a primary monocyte transduced with EGFP-CAAX (red) on hCMEC/D3 monolayer pre-stained with CellTrace (cyan) before the beginning of imaging. Z stacks images were taken every 10 min for 9 h and 50 min. Scale bar: 10 μm.

**Movie S4. Live cell imaging of an EGFP-OCLN-ΔC-expressing primary monocyte.** 3D time lapse spinning disk confocal imaging of a primary monocyte transduced with EGFP-OCLN-ΔC (red) on hCMEC/D3 monolayer pre-stained with CellTrace (cyan) before the beginning of imaging. Z stacks images were taken every 10 min for 6 h and 40 min. Scale bar: 10 μm.

**Movie S5. Live cell imaging of HIV-carrying monocytes infiltrating cortical organoids.** Epifluorescence Time-lapse imaging following protocol showed in Figure 5A. Primary human monocytes were pre-stained with CellTrace (cyan), exposed to HIV-1 Gag-imCherry (magenta) for 6 h prior the beginning of co-incubation of monocytes with a hESC-derived cortical organoid. Images were taken every hour for 8 days on a Nikon Biostation in BSL-3 environment.

## References

1. Terry RL, Getts DR, Deffrasnes C, van Vreden C, Campbell IL, King NJ. Inflammatory monocytes and the pathogenesis of viral encephalitis. J Neuroinflammation. 2012;9:270.

2. Garre JM, Yang G. Contributions of monocytes to nervous system disorders. J Mol Med (Berl). 2018;96(9):873–83.

3. Gerhardt T, Ley K. Monocyte trafficking across the vessel wall. Cardiovasc Res. 2015;107(3):321–30.

4. Gaudin R, Brychka D, Sips GJ, Ayala-Nunez V. Targeting tight junctions to fight against viral neuroinvasion. Trends Mol Med. 2022;28(1):12–24.

5. Saylor D, Dickens AM, Sacktor N, Haughey N, Slusher B, Pletnikov M, et al. HIV-associated neurocognitive disorder--pathogenesis and prospects for treatment. Nat Rev Neurol. 2016;12(4):234–48.

6. Hazleton JE, Berman JW, Eugenin EA. Novel mechanisms of central nervous system damage in HIV infection. HIV AIDS (Auckl). 2010;2:39–49.

7. Ayala-Nunez NV, Gaudin R. A viral journey to the brain: Current considerations and future developments. PLoS Pathog. 2020;16(5):e1008434.

8. Churchill MJ, Gorry PR, Cowley D, Lal L, Sonza S, Purcell DF, et al. Use of laser capture microdissection to detect integrated HIV-1 DNA in macrophages and astrocytes from autopsy brain tissues. Journal of neurovirology. 2006;12(2):146–52.

9. Pino M, Erkizia I, Benet S, Erikson E, Fernandez-Figueras MT, Guerrero D, et al. HIV-1 immune activation induces Siglec-1 expression and enhances viral trans-infection in blood and tissue myeloid cells. Retrovirology. 2015;12:37.

10. Lambotte O, Taoufik Y, de Goer MG, Wallon C, Goujard C, Delfraissy JF. Detection of infectious HIV in circulating monocytes from patients on prolonged highly active antiretroviral therapy. J Acquir Immune Defic Syndr. 2000;23(2):114–9.

11. Sonza S, Mutimer HP, Oelrichs R, Jardine D, Harvey K, Dunne A, et al. Monocytes harbour replication-competent, non-latent HIV-1 in patients on highly active antiretroviral therapy. AIDS. 2001;15(1):17–22.

12. Zhu T. HIV-1 in peripheral blood monocytes: an underrated viral source. J Antimicrob Chemother. 2002;50(3):309–11.

13. Massanella M, Bakeman W, Sithinamsuwan P, Fletcher JLK, Chomchey N, Tipsuk S, et al. Infrequent HIV Infection of Circulating Monocytes during Antiretroviral Therapy. J Virol. 2019;94(1).

14. Vestweber D. How leukocytes cross the vascular endothelium. Nat Rev Immunol. 2015;15(11):692–704.

15. Edgar R, Domrachev M, Lash AE. Gene Expression Omnibus: NCBI gene expression and hybridization array data repository. Nucleic Acids Res. 2002;30(1):207–10.

16. Weksler BB, Subileau EA, Perriere N, Charneau P, Holloway K, Leveque M, et al. Blood-brain barrier-specific properties of a human adult brain endothelial cell line. FASEB J. 2005;19(13):1872–4.

17. Prior IA, Hancock JF. Compartmentalization of Ras proteins. J Cell Sci. 2001;114(Pt 9):1603–8.

18. Berger G, Goujon C, Darlix JL, Cimarelli A. SIVMAC Vpx improves the transduction of dendritic cells with nonintegrative HIV-1-derived vectors. Gene Ther. 2009;16(1):159–63.

19. Nusrat A, Brown GT, Tom J, Drake A, Bui TT, Quan C, et al. Multiple protein interactions involving proposed extracellular loop domains of the tight junction protein occludin. Mol Biol Cell. 2005;16(4):1725–34.

20. Lacaz-Vieira F, Jaeger MM, Farshori P, Kachar B. Small synthetic peptides homologous to segments of the first external loop of occludin impair tight junction resealing. J Membr Biol. 1999;168(3):289–97.

21. Tavelin S, Hashimoto K, Malkinson J, Lazorova L, Toth I, Artursson P. A new principle for tight junction modulation based on occludin peptides. Mol Pharmacol. 2003;64(6):1530–40.

22. Wong V, Gumbiner BM. A synthetic peptide corresponding to the extracellular domain of occludin perturbs the tight junction permeability barrier. J Cell Biol. 1997;136(2):399–409.

23. Chung NP, Mruk D, Mo MY, Lee WM, Cheng CY. A 22-amino acid synthetic peptide corresponding to the second extracellular loop of rat occludin perturbs the blood-testis barrier and disrupts spermatogenesis reversibly in vivo. Biol Reprod. 2001;65(5):1340–51.

24. Everett RS, Vanhook MK, Barozzi N, Toth I, Johnson LG. Specific modulation of airway epithelial tight junctions by apical application of an occludin peptide. Mol Pharmacol. 2006;69(2):492–500.

25. Ayala-Nunez NV, Follain G, Delalande F, Hirschler A, Partiot E, Hale GL, et al. Zika virus enhances monocyte adhesion and transmigration favoring viral dissemination to neural cells. Nat Commun. 2019;10(1):4430.

26. Chang JM, Di Tommaso P, Taly JF, Notredame C. Accurate multiple sequence alignment of transmembrane proteins with PSI-Coffee. BMC Bioinformatics. 2012;13 Suppl 4(Suppl 4):S1.

27. Philip AM, Wang Y, Mauro A, El-Rass S, Marshall JC, Lee WL, et al. Development of a zebrafish sepsis model for high-throughput drug discovery. Mol Med. 2017;23:134–48.

28. Williams DW, Eugenin EA, Calderon TM, Berman JW. Monocyte maturation, HIV susceptibility, and transmigration across the blood brain barrier are critical in HIV neuropathogenesis. J Leukoc Biol. 2012;91(3):401–15.

29. Berre S, Gaudin R, Cunha de Alencar B, Desdouits M, Chabaud M, Naffakh N, et al. CD36-specific antibodies block release of HIV-1 from infected primary macrophages and its transmission to T cells. J Exp Med. 2013;210(12):2523–38.

30. Smith MS, Bentz GL, Alexander JS, Yurochko AD. Human cytomegalovirus induces monocyte differentiation and migration as a strategy for dissemination and persistence. J Virol. 2004;78(9):4444–53.

31. Du D, Xu F, Yu L, Zhang C, Lu X, Yuan H, et al. The tight junction protein, occludin, regulates the directional migration of epithelial cells. Dev Cell. 2010;18(1):52–63.

32. Sandig M, Korvemaker ML, Ionescu CV, Negrou E, Rogers KA. Transendothelial migration of monocytes in rat aorta: distribution of F-actin, alpha-catnin, LFA-1, and PECAM-1. Biotech Histochem. 1999;74(6):276–93.

33. McArdle S, Chodaczek G, Ray N, Ley K. Intravital live cell triggered imaging system reveals monocyte patrolling and macrophage migration in atherosclerotic arteries. J Biomed Opt. 2015;20(2):26005.

34. Rescigno M, Urbano M, Valzasina B, Francolini M, Rotta G, Bonasio R, et al. Dendritic cells express tight junction proteins and penetrate gut epithelial monolayers to sample bacteria. Nat Immunol. 2001;2(4):361–7.

35. Blank F, Wehrli M, Lehmann A, Baum O, Gehr P, von Garnier C, et al. Macrophages and dendritic cells express tight junction proteins and exchange particles in an in vitro model of the human airway wall. Immunobiology. 2011;216(1-2):86–95.

36. Muller WA. Mechanisms of leukocyte transendothelial migration. Annu Rev Pathol. 2011;6:323–44.

37. Morimoto S, Nishimura N, Terai T, Manabe S, Yamamoto Y, Shinahara W, et al. Rab13 mediates the continuous endocytic recycling of occludin to the cell surface. J Biol Chem. 2005;280(3):2220–8.

38. Shimizu Y, Shirasago Y, Suzuki T, Hata T, Kondoh M, Hanada K, et al. Characterization of monoclonal antibodies recognizing each extracellular loop domain of occludin. J Biochem. 2019.

39. Ploss A, Evans MJ, Gaysinskaya VA, Panis M, You H, de Jong YP, et al. Human occludin is a hepatitis C virus entry factor required for infection of mouse cells. Nature. 2009;457(7231):882–6.

40. Deffieu MS, Clement CMH, Dorobantu CM, Partiot E, Bare Y, Faklaris O, et al. Occludin stalls HCV particle dynamics apart from hepatocyte tight junctions, promoting virion internalization. Hepatology. 2022;76(4):1164–79.

41. Paul CD, Devine A, Bishop K, Xu Q, Wulftange WJ, Burr H, et al. Human macrophages survive and adopt activated genotypes in living zebrafish. Sci Rep. 2019;9(1):1759.

42. Honeycutt JB, Wahl A, Baker C, Spagnuolo RA, Foster J, Zakharova O, et al. Macrophages sustain HIV replication in vivo independently of T cells. J Clin Invest. 2016;126(4):1353–66.

43. Honeycutt JB, Thayer WO, Baker CE, Ribeiro RM, Lada SM, Cao Y, et al. HIV persistence in tissue macrophages of humanized myeloid-only mice during antiretroviral therapy. Nat Med. 2017;23(5):638–43.

44. Ganor Y, Real F, Sennepin A, Dutertre CA, Prevedel L, Xu L, et al. HIV-1 reservoirs in urethral macrophages of patients under suppressive antiretroviral therapy. Nat Microbiol. 2019;4(4):633–44.

45. Wong ME, Jaworowski A, Hearps AC. The HIV Reservoir in Monocytes and Macrophages. Front Immunol. 2019;10:1435.

46. Igarashi T, Brown CR, Endo Y, Buckler-White A, Plishka R, Bischofberger N, et al. Macrophage are the principal reservoir and sustain high virus loads in rhesus macaques after the depletion of CD4+ T cells by a highly pathogenic simian immunodeficiency virus/HIV type 1 chimera (SHIV): Implications for HIV-1 infections of humans. Proceedings of the National Academy of Sciences of the United States of America. 2001;98(2):658–63.

47. Gupta RK, Abdul-Jawad S, McCoy LE, Mok HP, Peppa D, Salgado M, et al. HIV-1 remission following CCR5Delta32/Delta32 haematopoietic stem-cell transplantation. Nature. 2019;568(7751):244-8.

48. Henrich TJ, Hanhauser E, Marty FM, Sirignano MN, Keating S, Lee TH, et al. Antiretroviral-free HIV-1 remission and viral rebound after allogeneic stem cell transplantation: report of 2 cases. Annals of internal medicine. 2014;161(5):319–27.

49. Cummins NW, Rizza S, Litzow MR, Hua S, Lee GQ, Einkauf K, et al. Extensive virologic and immunologic characterization in an HIV-infected individual following allogeneic stem cell transplant and analytic cessation of antiretroviral therapy: A case study. PLoS Med. 2017;14(11):e1002461.

50. Veenstra M, Leon-Rivera R, Li M, Gama L, Clements JE, Berman JW. Mechanisms of CNS Viral Seeding by HIV(+) CD14(+) CD16(+) Monocytes: Establishment and Reseeding of Viral Reservoirs Contributing to HIV-Associated Neurocognitive Disorders. mBio. 2017;8(5).

51. Morner A, Bjorndal A, Albert J, Kewalramani VN, Littman DR, Inoue R, et al. Primary human immunodeficiency virus type 2 (HIV-2) isolates, like HIV-1 isolates, frequently use CCR5 but show promiscuity in coreceptor usage. J Virol. 1999;73(3):2343–9.

52. Gaudin R, Berre S, Cunha de Alencar B, Decalf J, Schindler M, Gobert FX, et al. Dynamics of HIV-containing compartments in macrophages reveal sequestration of virions and transient surface connections. PLoS One. 2013;8(7):e69450.

